# Foxe1 orchestrates thyroid and lung cell lineage divergence in mouse stem cell-derived organoids

**DOI:** 10.1101/2022.05.16.492074

**Authors:** Barbara F. Fonseca, Cindy Barbée, Mirian Romitti, Sema Elif Eski, Pierre Gillotay, Daniel Monteyne, David Perez Morga, Samuel Refetoff, Sumeet Pal Singh, Sabine Costagliola

## Abstract

Patterning of endoderm into lung and thyroid lineages depends upon a correct early expression of a homeobox domain-containing transcription factor, Nkx2-1. However, the gene networks distinguishing the differentiation of those lineages remain largely unknown. In the present work, by using mouse embryonic stem cell lines, single-cell RNA sequencing, and transcriptomic and chromatin accessibility profiling, we show that knockout of Foxe1 drastically impairs Nkx2-1+ cells differentiation and maturation into thyroid follicular-like cells. Concomitantly, a subset of Foxe1 null/Nkx2-1+ cells have a remarkable ability *in vitro* to undergo a lung epithelial differentiation program and form lung-like organoids harboring cells transcriptionally similar with mouse fetal airway and alveolar cell types. These results demonstrate, for the first time, lung lineage derivation at the expense of thyroid lineage, by a simple removal of a transcription factor, and provide insights into the intricated mechanisms of fate decisions of endodermal cell types.

**Highlights:**

- Forward programming of mESCs with transient Nkx2-1 and Pax8 overexpression, followed by c-AMP treatment, leads to differentiation of functional thyroid follicles *in vitro*;
- In absence of Foxe1, thyroid follicle-like structures, derived from mESCs, are scarce and non-functional;
- Concomitantly, a subset of Nkx2-1-expressing cells generated from Foxe1KO mESCs spontaneously form lung organoids containing multiple differentiated lung cell types;
- ATACseq analyses show higher chromatin remodeling in Nkx2-1-expressing cells in control compared to Foxe1KO cells, especially for genes involved in thyroid maturation and maintenance of the 3D structure of the follicle.

## Introduction

Embryonic development comprises a stepwise progression towards restriction of cell potentiality and acquisition of a differentiated cellular state. There are various molecular mechanisms controlling cell fate commitment, involving cellular response to extracellular cues, chromatin remodeling and transcription factors activation of lineage-related cis-regulatory regions. Unveiling the molecular events triggering lineage decisions is not only crucial to understand the biology behind those phenomena but also to allow *in vitro* cell engineering, which is a potential avenue for disease modeling and regenerative applications in medicine.

During endoderm organogenesis, the space and time coordinated expression of transcription factors such as Sox2, Cdx, Foxa2, Hhex, Pdx1, among others, patterns the endoderm layer into more specific cell lineages (Kraus and Grapin-Botton, 2012; Zorn and Wells, 2009). For thyroid and lung derivation, for example, it is broadly known that the homeodomain-containing factor *Nkx2-1* (also called thyroid transcription factor 1; *Ttf-1*), is the first gene for which expression is detected in specific domains of the ventral anterior foregut endoderm, where the thyroid and lung primordia arise around embryonic days 8-9 during mouse embryogenesis (Cardoso and Lu, 2006; Lazzaro et al., 1991). In addition, *Nkx2-1* expression is observed throughout lung and thyroid embryonic and adult life. Therefore, it is not surprising that NKX2-1 gene mutations, in humans and mice, engender a variety of thyroid and pulmonary abnormalities, as well as neurological defects (Butt et al., 2008; Herriges and Morrisey, 2014; Willemsen et al., 2005).

Besides the requirement of Nkx2-1 signaling for both thyroid and lung lineages, the molecular pathways leading to their specification are still poorly understood. For this reason, in early directed differentiation protocols to derive thyroid/lung lineages from mouse embryonic stem cells (mESC), a dual generation of thyroid/lung Nkx2-1 progenitor cells was obtained (Longmire et al., 2012). The better understanding of the different requirements for Fgf and Wnt pathways activation was key to separately drive each lineage in mESCs, after an *in vitro* step of anterior foregut specification (Dame et al., 2017; Kurmann et al., 2015; Mou et al., 2012; Serra et al., 2017). More recent reports are helping to shed light on the early stages of thyroid and lung organogenesis (Haerlingen et al., 2019; Ikonomou et al., 2020; Rankin et al., 2021; Vandernoot et al., 2021). Overall, those data indicate an intricate relationship among thyroid/lung lineages, at early stages of mammalian development.

At the time of thyroid specification, besides Nkx2-1, cells from thyroid anlage are identified by the restricted expression of three other transcription factors: Pax8, Hhex and Foxe1 (López-Márquez et al., 2021). According to proposed mechanisms for thyroid specification (Parlato et al., 2004), the onset of Foxe1 expression is downstream to those cited players and Foxe1 (also called thyroid transcription factor 2; Ttf-2) is key to the induction of more thyroid-restricted markers, such as *Thyroglobulin* and *Thyroperoxidase* (*Tpo)* (Aza-Blanc et al., 1993; Francis-Lang et al., 1992; López-Márquez et al., 2019; Santisteban et al., 1992). Foxe1 is a member of the Forkhead family, a group of transcription factors classically involved in endoderm lineage decisions, acting directly to induce the transcription program of a particular cell lineage but also potentially repressing alternative cell fates (Golson and Kaestner, 2016; Li et al., 2016; Sekiya and Suzuki, 2011; Zaret and Carroll, 2011). Interestingly, at the stage of thyroid/lung lineages commitment in the anterior foregut endoderm, *Foxe1* is expressed throughout the anterior foregut domain but it is specifically absent in the lung primordium (De Felice and Di Lauro, 2004; Kuwahara et al., 2020), suggesting that *Foxe1* expression in this region is not compatible with the correct lung lineage establishment.

In the present work, we investigated the role of Foxe1 in the differentiation of thyroid lineage and asked whether Foxe1 absence can influence lineage decision of Nkx2-1+ progenitors *in vitro*. For this, we used our mESC-based model, in which thyroid differentiation is induced by Nkx2-1 and Pax8 transient overexpression and formation of functional thyroid follicle-like structures is obtained by recombinant TSH treatment (Antonica et al., 2012; Romitti et al., 2021). Here we show that, without Foxe1, Nkx2-1+ precursors fail to form polarized and mature thyroid follicular cells whereas a large subset of Nkx2-1+ cells are able to diverge to an alternative lineage pathway, giving rise to multiple lung cell types *in vitro*.

## Results

### Foxe1 is required for thyroid development *in vitro*

Seminal works have previously shown that Foxe1 depletion/misfunction in the thyroid gland results in various thyroid defects, from the deregulation of key players involved in thyroid hormone production pathway up to the agenesia of the gland (Clifton-Bligh et al., 1998; De Felice et al., 1998; Fernández et al., 2013). To further investigate the role of Foxe1 in thyroid follicular cells and to validate our mESC-derived thyroid organoid model to study genes causing hypothyroidism, we mutated Foxe1 loci in mESCs derived from our previously established transgenic line (A2lox-Nkx2-1- Pax8 line). In this line, transient overexpression of Nkx2-1 and Pax8, followed by 2- weeks treatment with hTSH/ or c-AMP analogues, yields self-organized thyroid follicles that secrete T4 hormone *in vitro* with high efficiency (Antonica et al., 2012) (Figure 1A-B). In addition, differentiation efficiency can be visually assessed using an EGFP transgene inserted into the hypoxanthine phosphoribosyltransferase (HPRT) locus of mESC lines under the control of a bovine-responsive thyroglobulin (Tg) promoter (Romitti et al., 2021).

**Figure 1.**
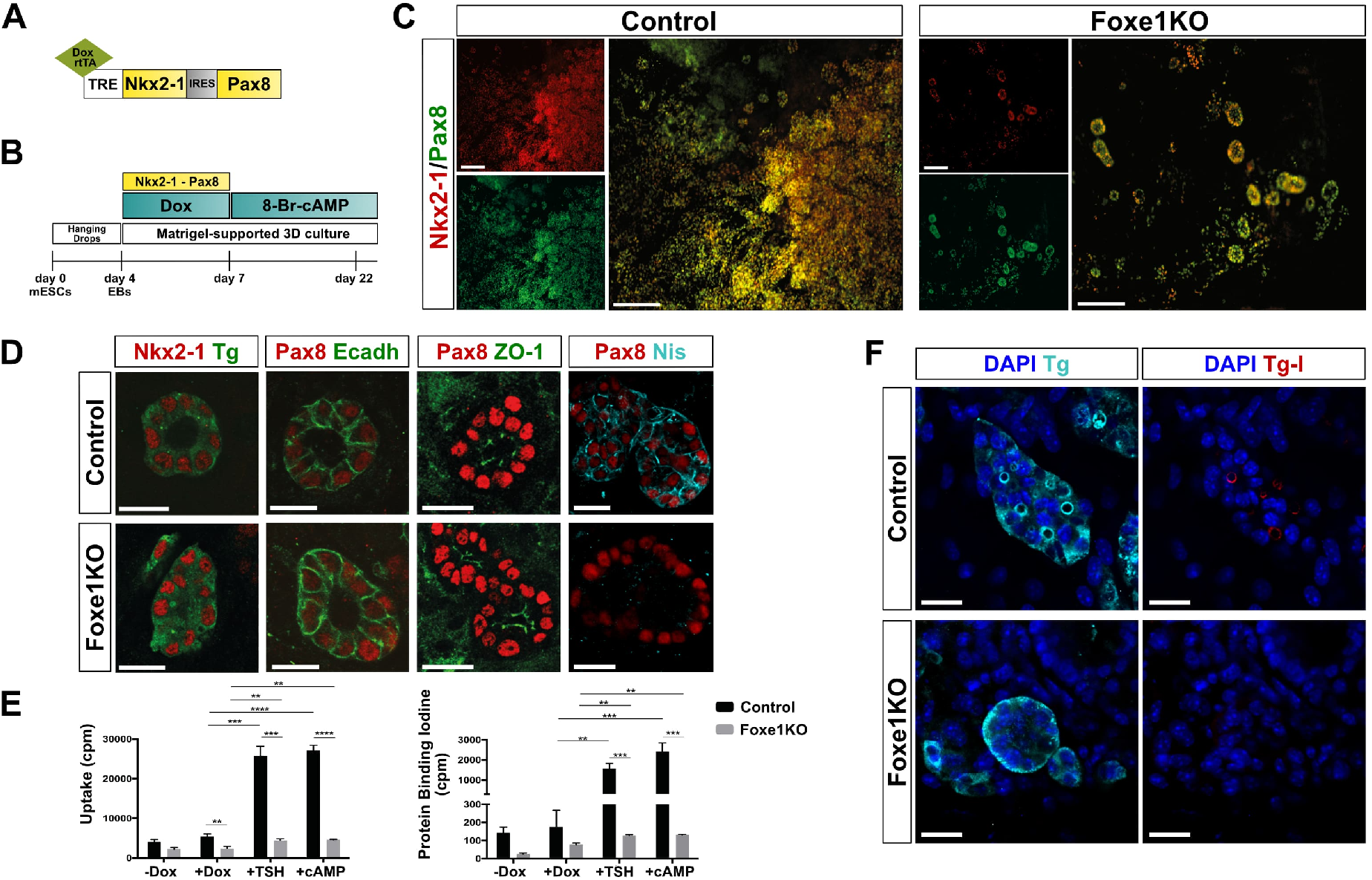
Foxe1 gene is required for efficient thyroid differentiation *in vitro*. (**A**) Schematic representation of tetracycline-inducible cassette to drive co-expression of Nkx2-1 and Pax8. (**B**), Diagram of the 22-days differentiation protocol for mESCs. (**C-F**) Immunostaining and organification assays performed at day 22 of differentiation. (**C**) Immunostaining of double positive Nkx2-1 and Pax8 thyroid follicles in control and Foxe1KO cells. (**D, E**) Immunostaining of a panel of thyroid markers in control and Foxe1KO cells (**F**) Iodide uptake and organification assays. Histograms represent the radioactivity (Uptake-cpm) of the cell iodine-125 uptake (left) and the radioactivity of protein-bound [^125^I] (PBI-cpm) measured using a γ-counter (right). Unpaired t-test. *P<0,05, **P<0,01, ***P<0,001. Scale bars: 150 µm (**C**); 20 µm (**D, E**). Tg: thyroglobulin, Ecadh: Ecadherin, Tg-I: iodinated thyroglobulin.

To obtain Foxe1 knockout (KO) lines, Foxe1 loci were edited using TALEN technology and clones selected contained mutations that resulted in a premature stop codon (Figure S1A). After validation of the Foxe1KO lines with respect to maintenance of mESC pluripotency and responsiveness to the Nkx2-1/Pax8 tetracycline-inducible cassette (Figure S1B-C), control and Foxe1KO lines were subjected to the differentiation protocol (Figure 1B). Three days after doxycycline treatment (day 7 of differentiation protocol), control and Foxe1KO lines are similarly efficient to activate exogenous expression of the artificial Nkx2-1/Pax8 transgenes (Figure S2A). This exogenous transgene expression leads to the activation of endogenous Nkx2-1 and Pax8 loci (Antonica et al., 2012). All lines successfully achieved overexpression of the endogenous loci of Nkx2-1 and Pax8. In contrast, the expression of Hhex and Foxe1 was drastically decreased in Foxe1KO cells (Figure S2A).

At the end of the differentiation protocol (Figure 1B), the effect of Foxe1 depletion during thyroid formation *in vitro* was clearly observed. First, we detected, by RT-qPCR, a strong reduction in mRNA levels of many thyroid-related genes (Figure S2B). Moreover, the number of thyroid follicles formed in the absence of Foxe1 was greatly reduced compared with the control line (Figure 1C). Although the formation of scarce one-cell layer follicular structures could still be detected, protein expression of transporters essential for thyroid maturation and function, such as the sodium/iodide symporter Nis (Slc5a5), was drastically reduced (Figure 1D). Finally, thyroid follicles from the control line displayed proper iodination of Tg (Figure 1E) and high uptake of radioactive iodide and iodide binding to Tg, whereas these processes were absent in Foxe1-depleted cells (Figure 1F). Because Tshr mRNA levels were drastically low in the absence of Foxe1 (Figure S2B), we replaced the 2-weeks recombinant hTSH treatment with a c- AMP analogue, as TSH ligands signal through the cAMP pathway (Kimura, 2001). The cAMP treatment produced similar results regarding thyroid-related gene expression and iodide organification *in vitro* (Figure 1F and S2B). Overall, these results demonstrate the drastic effect of Foxe1 knockout on thyroid lineage derivation *in vitro* and validate our mESC-model to investigate genes involved in thyroid dysgenesis.

### Absence of Foxe1 promotes lung differentiation *in vitro*

Besides the impairment of thyroid generation in Foxe1KO mESCs, we observed an additional, albeit striking, phenotype: the self-assembly of Nkx2-1-expressing cells into epithelial structures with a morphology clearly different from thyroid follicles, in all culture wells of Foxe1KO samples (Figure 2A). Because Nkx2-1 is involved in the specification of another ventral foregut derivative, the lung (Cardoso and Lu, 2006; Lazzaro et al., 1991), we wondered whether the appearance of these 3D structures might indicate spontaneous generation of lung tissue in culture. By immunostaining analyses at day 22, we observed that these Nkx2-1+ organoids indeed expressed typical lung-related markers, such as Trp63/Krt5 (basal cells), Hopx (alveolar cells), Muc5ac (goblet cells) and Scgb3a2 (club cells) (Figure 2B-E). In addition, upregulation of lung-related transcripts was observed in Foxe1KO cells, as detected by RT-qPCR (Figure S3A). Ultrastructural microscopy of these Foxe1KO-derived organoids revealed the presence of bronchiole-like structures and cells exhibiting the morphology of typical lung cell types, such as multiciliated cells, mucus-secreting cells (goblet cells), and type 1 and 2 alveolar cells (Figure 2F and S3B-M). Altogether, these results suggest that branched lung structures are formed from Foxe1-depleted mESCs, extending from airway components to the alveolar-like cells.

**Figure 2.**
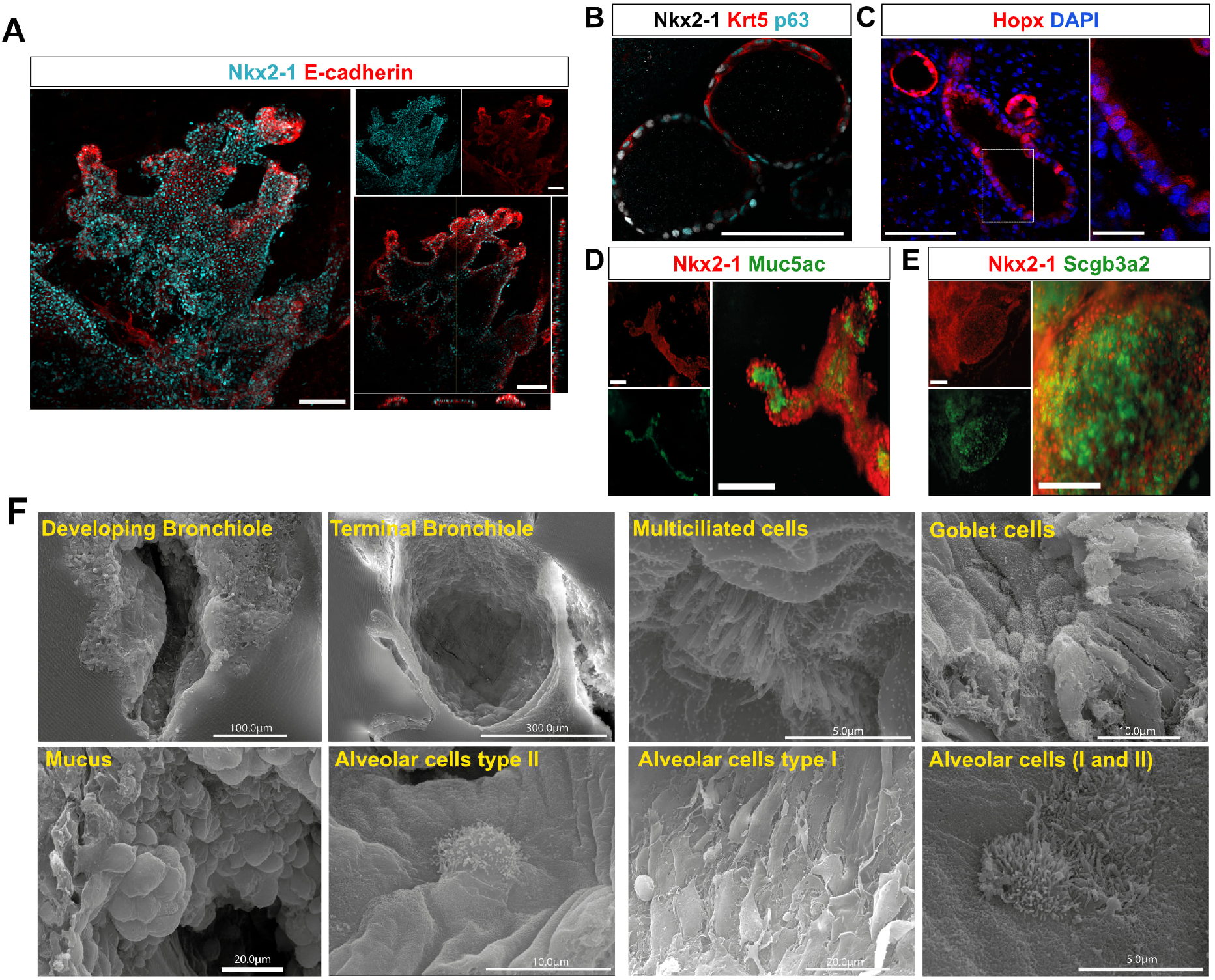
Generation of pulmonary structures in absence of Foxe1. (**A**) Orthogonal view by confocal microscopy of *in vitro* lung-like structures identified by immunostaining of Nkx2-1 and E-cadherin at day 22 of differentiation protocol. (**B-E**) Immunostaining of Foxe1KO mESC-derived organoids with a panel of lung-related markers: basal cells (Nkx2-1, Krt5 and Trp63); alveolar cells (Hopx); goblet cells (Nkx2-1 and Muc5ac) and secretory (club) cells (Nkx2-1 and Scgb3a2). (**F**) Scanning electron microscopy at day 22 Foxe1KO mESC-derived pulmonary structures: a large panel of pulmonary cell lineages are evidenced, including airway and alveolar cells. Scale bars : 100µm (**A**); 25µm (**B**); 100µm (**C**; left) and 25µm (**C**; right); 150µm (**D,E**).

### Characterization of Foxe1KO-derived thyroid and lung cell populations at the single-cell resolution

The observation that Foxe1 depletion in mESCs results in a decreased thyroid cells generation and the appearance of multiple lung-related cell types, prompted us to better characterize the cell population derived from Foxe1KO cells, by scRNAseq. For this, we first designed Nkx2-1 reporter lines, allowing easier identification and isolation of thyroid and lung cell types. A mKO2 fluorescent tag was inserted into the 3’ end of the endogenous Nkx2-1 loci of the control A2lox-Nkx2-1-Pax8 line (Figure S4A-C). The reporter fluorescence could be visualized by microscopy and flow cytometry (Figure S4D-E). Foxe1KO/Nkx2-1 reporter lines were created and differentiated using our differentiation protocol (Figure S4F-I). The same Foxe1KO phenotype (i.e., disruption of thyroid function and appearance of lung-like organoids) was observed in these new lines (Figure S5).

For the single cell profiling of Foxe1KO cells, we enriched by FACS three cell populations: Nkx2-1/mKO2+/bv Tg prom/EGFP+ (15%), Nkx2-1/mKO2+/bv Tg prom/EGFP- (50 %) and Nkx2-1/mKO2- (35%) (Figure 3A and S6A). A total of 12000 cells were profiled for scRNA-seq using the droplet-based assay from 10xGenomics system. After quality control, we obtained 7523 cells. The profiled cells could be divided into 12 clusters composed of Nkx2-1+ cells (i.e. Thyroid1, Thyroid2, Lung1, Lung2, Nkx2-1+Pax8- 1, Nkx2-1+Pax8- 2) and Nkx2-1 negative cells (i.e. Epithelial cells, Prolif. epithelial cells, Basal Epithelium, Pluripotent cells, Mesenchymal cells) (Figure 3B). Gene Ontology and literature mining were used to characterize the clusters based on top differentially expressed genes (Figure S6B-C). Among Nkx2-1 negative cells, epithelial-like cells are found in three clusters (702 broad Epithelial cells, 244 Prolif. Epithelial cells and 123 Basal epithelial cells). 1411 pan-mesenchymal cells (Mesenchymal cells cluster), 280 cells expressing endoderm markers (Endoderm cluster) and a remaining pluripotent cell population were also found (Pluripotent cells cluster, 520 cells).

**Figure 3:**
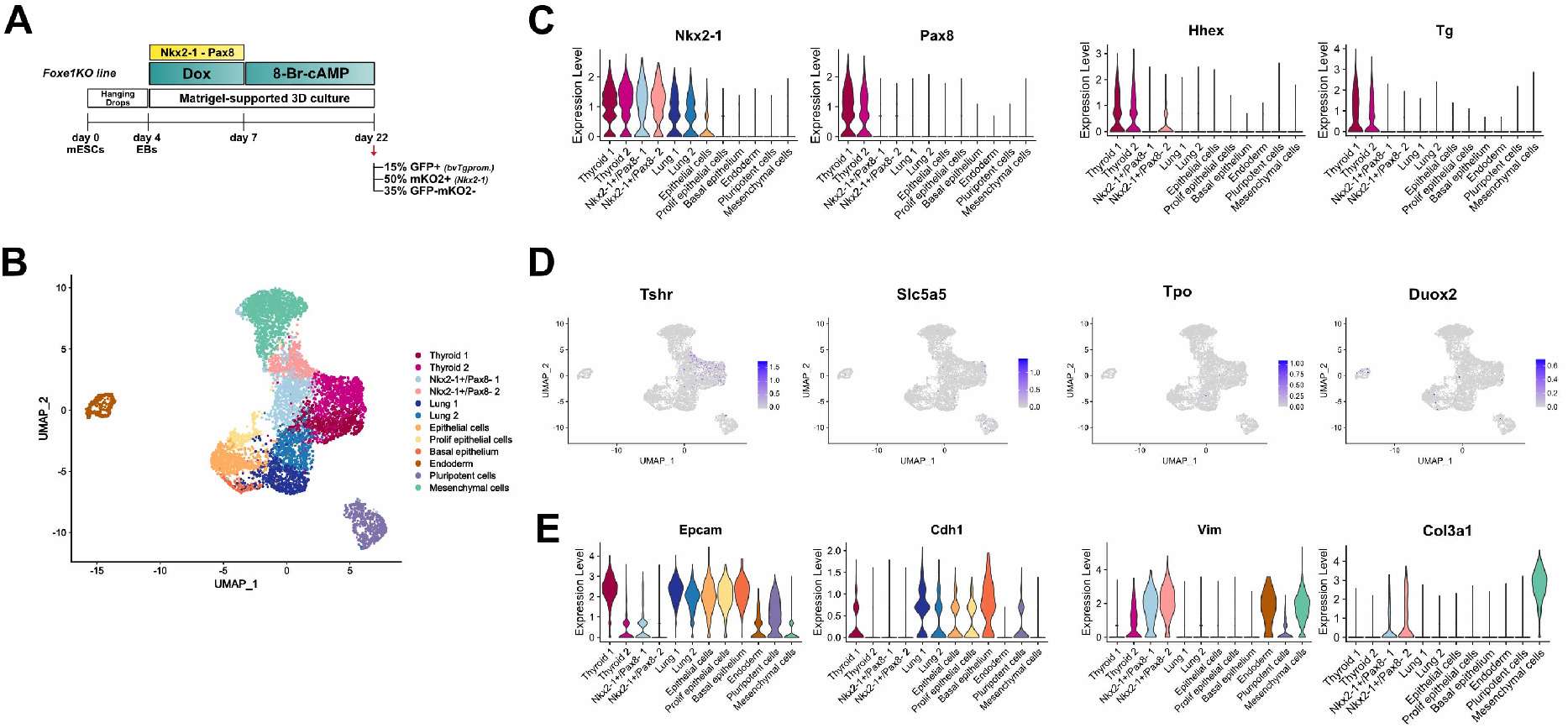
Single-cell RNA-seq analysis of Foxe1KO-derived thyroid cells. (**A**) Schematic diagram of differentiation protocol and purification of distinct cell populations at day 22 for scRNA-seq experiment. (**B**) Unsupervised clustering of 7523 single-cells profiled, colored by cluster assignment. (**C**) Violin plots featuring the average expression per cluster of early thyroid markers. (**D**) UMAP expression plots of markers for thyroid maturation (**E**) Violin plots featuring average expression per cluster of selected epithelial and mesenchymal markers.

### Single cell RNA-seq analysis reveals that Foxe1KO cells lack functional thyroid transcripts and evidences aberrant Nkx2-1+ cells

Six clusters in Foxe1KO sample consist of cells expressing Nkx2-1. Thyroid 1 (722 cells) and Thyroid 2 (947 cells) clusters exhibit a typical thyroid signature characterized by the expression of *Pax8*, *Hhex*, and *Tg* (Figure 3B-C and SB-D). Note that expression of mature markers such as *Tpo*, *Slc5a5*, and *Duox2* is completely absent, while *Tshr* levels are low, supporting our previous results (Figure 3D and S6B- D). Among those thyroid clusters, the normalized average expression of *Pax8*, *Hhex* and *Tg* appears to be lower in Thyroid 2 than in Thyroid 1 group. In addition, Thyroid 2 cell group lacks expression of *Epcam* and *Cdh1* (Figure 3E and S6B-D). Since E-cadherin expression is maintained throughout thyroid morphogenesis *in vivo* (Fagman et al., 2003), this finding suggests that Thyroid 2 cells are abnormal *in vitro*-derived thyrocytes not organized into follicular units.

We also identified two additional cell clusters characterized by strong *Nkx2-1* but absent *Pax8* expression (Nkx2-1+/Pax8- 1 and Nkx2-1+/Pax8- 2 clusters, with 843 and 489 cells, respectively). Their transcriptomic signature suggest that they are distinctly different from the thyroid and lung clusters. For example, they lack *Epcam* and *Cdh1* expression but are instead enriched in mesenchymal-like markers such as *Vim*, *Acta2* and *Col3a1* (Figure 3E). Gene Ontology analysis for these clusters show enrichment in the “extracellular matrix/epithelial to mesenchymal transition” and “glycolysis/hypoxia pathways”, respectively (Figure S6C). Since mesenchymal-like cells expressing Nkx2-1+ are not described during development, these cells likely reflect part of an *in vitro* Foxe1KO phenotype, along with the appearance of lung tissue *in vitro*. Interestingly, these cells do not express neural markers such as *Ascl1*, *Tubb3*, *Pax6* and *Six3*, making it unlikely that they possess a Nkx2-1+ neural signature. Finally, thyroid C cells also express *Nkx2-1* (Nilsson and Williams, 2016), but no co-expression of *Calca*, *Foxa1* and *Foxa2* was observed in these cells (data not shown).

### Foxe1KO mESCs differentiate into multiple lung cell types harboring transcriptomic signatures encountered in E17.5 embryonic mouse lung tissue

Lung-related cells originated from Foxe1KO cells are present in the Lung 1 and Lung 2 clusters (668 and 574 cells, respectively). To better define specific lung cell types, and avoid contamination with thyrocytes, we computationally selected cells expressing *Nkx2-1* and *Epcam*, but lacking *Tg* and *Pax8*, from the Foxe1KO dataset. After re-clustering and assigning the cell populations, we identified eight different lung subsets (Figure 4A). Cell types were identified using a signature score index based on the top 20 marker genes expressed by several lung epithelial cell types found in the scRNAseq dataset of E17.5 mouse lung tissue (Frank et al., 2019). Since this dataset includes lung epithelial cells derived from murine alveolar and terminal bronchioles, a literature mining was performed to define the signature score of basal cells found mainly in the upper airways (Table S1). As a result, we identified cell populations with a strong signature of secretory cells (cluster c), basal cells (cluster a), multiciliated cells (cluster h), and a small group with a slightly enriched signature for alveolar type 1 (AT1) cells (cluster g). In addition, we identified a cluster enriched in both basal and secretory markers (cluster d), representing cells transitioning from basal to secretory cell fate (Figure 4A-B). Finally, we detected two clusters, b and f, that were highly enriched in Sox9 and Igf1 transcripts, respectively (Figure 4B). These factors are highly expressed in early lung development, with Sox9 specifically present in distal bud cells (Nikolić et al., 2018).

**Figure 4:**
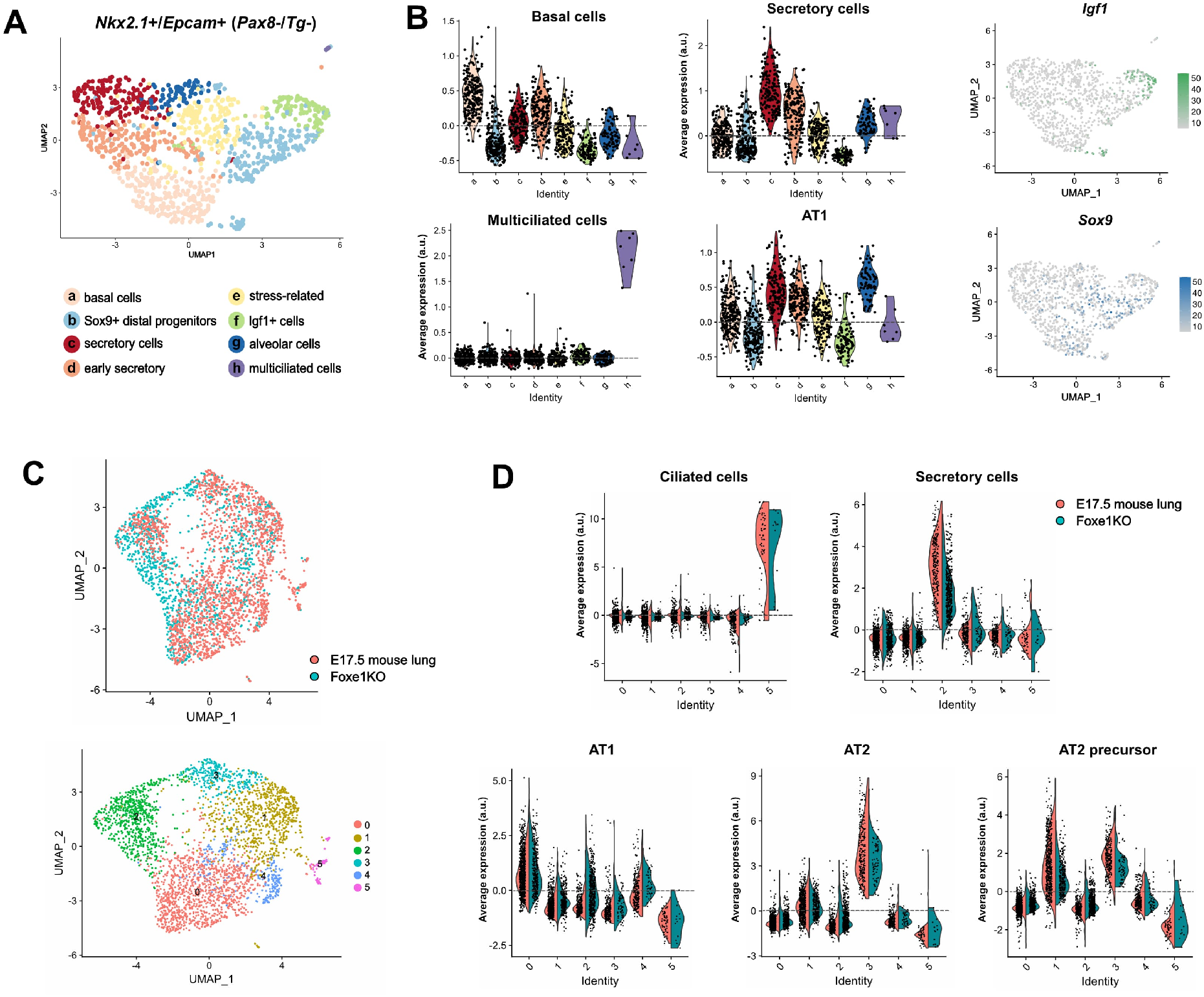
Single-cell RNA-seq analysis of Foxe1KO-derived lung cells. (**A**) 1155 cells co-expressing Nkx2-1 and Epcam (but not expressing Pax8 and Tg) were computationally isolated, re-clustered and cellular transcriptome heterogeneity visualized using UMAP. (**B**) Average expression levels per cluster of indicated lung cell type signatures and UMAP expression plot of Igf1 and Sox9 normalized expression, defining two clusters of early lung progenitors. (**C**) UMAP plot of Foxe1KO cells (1155) integrated with E17.5 mouse lung Nkx2-1+Epcam+ cells (2000 cells) (upper graph). 6 identified clusters contain both in vitro and in vivo derived lung cells (bottom graph). (**D**) Average expression levels of indicated lung cell type signatures identified in both mouse lung (pink) and Foxe1KO cells (blue).

With the purpose to better characterize Foxe1KO-derived lung cell types and to test the degree of similarity with in vivo encountered lung cells, we performed scRNAseq comparative analyses between the above cited E17.5 mouse lung dataset and Foxe1KO-derived lung cells (Frank et al., 2019; Stuart et al., 2019) (Figure 4C- D). To do this, we integrated *Nkx2-1*+/*Epcam*+/*Tg*-/*Pax8*- cells from the Foxe1KO dataset with *Nkx2-1*+/*Epcam*+ cells from the *in vivo* data. The analysis revealed a high overlap among cells from the two datasets (Figure 4C). Furthermore, when we tested for enriched signatures of multiple lung cell types based on the normalized average expression of marker genes, we detected the presence of alveolar cells (AT1, AT2, and AT2 precursors) in groups 0, 3 and 1, respectively, in both datasets (Figure 4D). Ciliated and secretory cells were also identified, as shown in Figure 4B. Of note, because the mouse fetal lung dataset is devoid of basal cells (Frank et al., 2019), this particular lung cell type could not be addressed with this integrated analysis.

In summary, Foxe1 depletion abrogates the proper differentiation of thyrocytes and instead allows the development of different lung cell types. In addition, the Foxe1KO-derived lung cells are remarkably similar to the cells encountered *in vivo* in terms of the transcriptomic signature of the major lung cell type markers.

### Chromatin accessibility analysis of Nkx2-1 expressing cells

Our results show that depletion of Foxe1 abrogates proper differentiation of Nkx2-1 cells into thyrocytes *in vitro* while leading to the appearance of lung cell types.

To identify potential molecular players involved in Foxe1 action, we performed a temporal analysis of the global chromatin accessibility of Nkx2-1-expressing cells, using bulk ATACseq. To develop an optimal experimental procedure, we first examined the kinetics of thyroid-related gene expression (e.g., *Nkx2-1*, *Foxe1* and *Tg*) at key points of the protocol in control and Foxe1KO cells (Figure 5A-D). After induction of the artificial Nkx2-1/Pax8 cassette between days 4 and 7, exogenous Nkx2-1 mRNA levels are dramatically reduced on day 8 and it is virtually abolished around day 11 (Figure 5A). Conversely, endogenous expression of Nkx2-1 increases rapidly as of day 7 and reaches a maximum at day 14 (Figure 5B). Foxe1 is firstly induced by the direct action of exogenous Nkx2-1 and Pax8 at day 7, following a down-regulation on days 7 and 9 (Figure5C). A second wave of induction occurs from day 9 onwards, likely due to the endogenous expression of Nkx2-1/Pax8, in conjunction with treatment with c-AMP (Antonica et al., 2012; Ortiz et al., 1997). As expected, Tg mRNA levels are very low in early stages of culture, but increase steadily after day 11, reaching the highest level at the end of the protocol (Figure 5D). The temporal expression of these thyroid markers is consistent with the expected pattern of thyroid differentiation genes (Fernández et al., 2015; Romitti et al., 2021). These results suggest that events important for thyroid specification *in vitro* occur before day 10, as high *Tg* levels are observed after this time point. Coincidentally, the increase in *Foxe1* expression occurs approximately one day before *Tg* expression.

**Figure 5:**
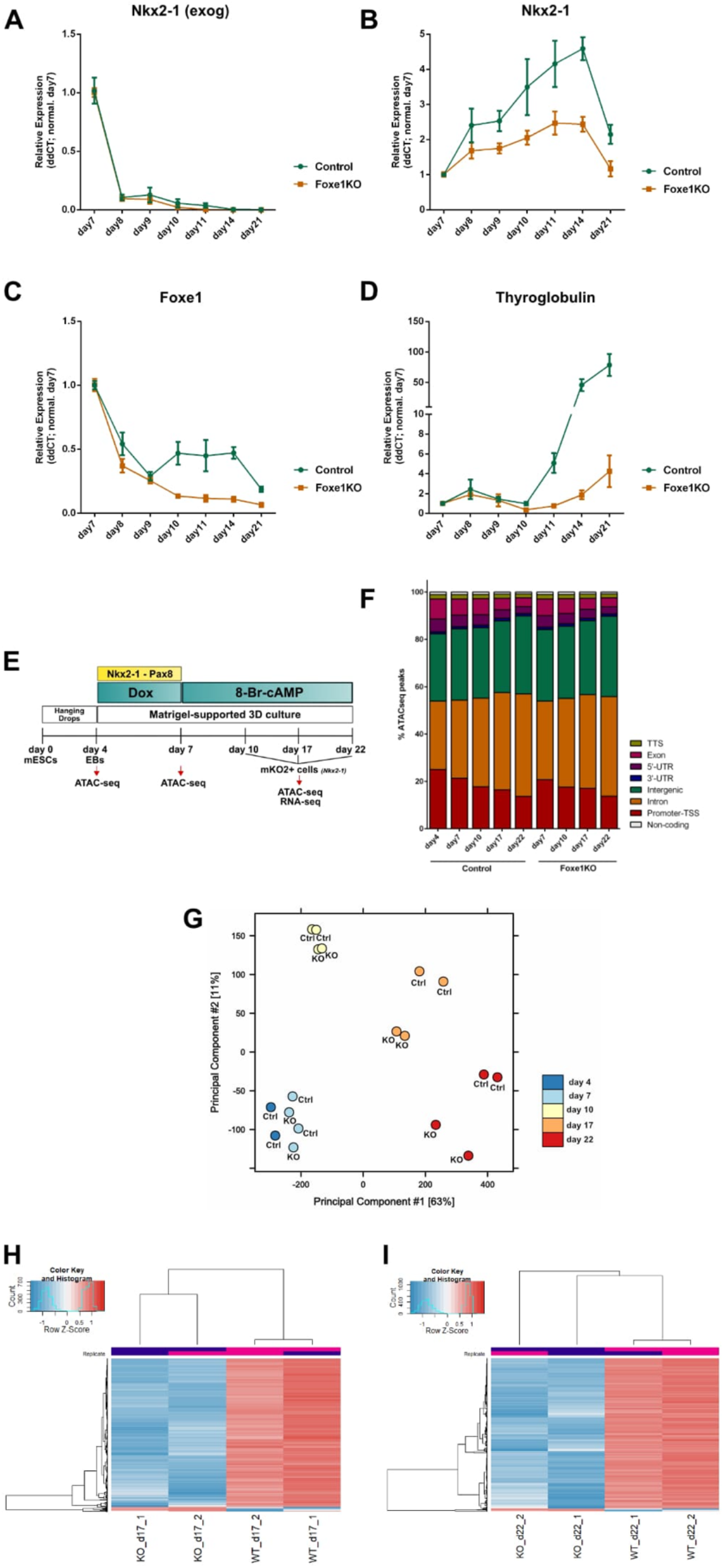
ATACseq profiling of control and Foxe1KO Nkx2-1 expressing cells at different stages of differentiation protocol. (**A-D**) Relative mRNA expression of thyroid marker genes through different time points of culture. Relative expression of each transcript is presented as fold change compared to cells from the first time point (day 7) as Mean +/- SEM. Exog: exogenous. (**E**) Schematic representation of differentiation protocol and time points of cell sorting of Nkx2-1+ (mKO2+) cells for ATAC-seq and RNA-seq analysis. (**F**) Total ATAC-seq peaks distribution in diverse genomic regions. (**G**) Principal component analysis (PCA) to compare total ATACseq peaks among time points. (**H-I**) Heatmaps of differentially accessible peaks between control and Foxe1KO at day 17 (**H**) and day 22 (**I**).

Based on the kinetics of expression of lineage-specific genes, we decided to isolate Nkx2-1+ cells in the control and Foxe1KO samples by FACS at day 10 to identify early progenitors of the thyroid and lung lineages, and more differentiated cell types at days 17 and 22. In addition, cells from day 4 and day 7, time points before and after doxycycline treatment, respectively, were also profiled (Figure 5E). In total, approximately 90,000 ATAC-seq peaks were obtained per sample, which were distributed in different open genomic regions (Figure 5F). PCA analysis suggests a gradual change in global chromatin accessibility that correlates with the timing of the differentiation protocol (Figure 5G). Day 4 and day 7 samples are relatively more similar to each other than the other late time points. In addition, no clear difference between the control and Foxe1KO is observed at day 7. Since the time point at day 7 corresponds to the end of doxycycline treatment, this implies that exogenous induction of Nkx2-1 and Pax8 results in a similar open chromatin landscape in both lines. ATACseq peaks originating from Nkx2-1+ cells (day 10, day 17 and day 22) show a gradual chromatin remodeling after day 10, which becomes more evident at days 17 and 22 (Figure 5G-I).

### Enrichment in thyroid maturation markers in control Nkx2-1+ cells and the identification of predicted Foxe1 target genes

Our results suggest that the most striking differences in chromatin accessibility between control and Foxe1KO are seen towards the end of the protocol (Figure 5G- I). Using the Diffbind package for differential binding analyses between control and Foxe1KO conditions, we identified more than 23547 genomic regions that are differentially accessible at day 17 (19112 are up-regulated in control in comparison with Foxe1KO and 4435 up in Foxe1KO versus control) (Figure 5H) and 14293 peaks at day 22 (13500 up-regulated in control in comparison with Foxe1KO and 793 up in Foxe1KO versus control) (Figure 5I). The number of opened chromatin regions in control cells is 17-fold greater than in Foxe1KO cells at day 22, indicating an important role for Foxe1 in opening specific chromatin domains that are otherwise silenced. Of note, more than 60% of the up-regulated transcripts found in bulk RNAseq analyses of the control line were associated with significant chromatin opening compared with Foxe1KO (Figure 6A). Among them are most of the genes associated with the thyroid gland, including essential genes for gland maturation and function, such as *Tshr*, *Tpo* and *Duox2* (Figure 6B).

**Figure 6:**
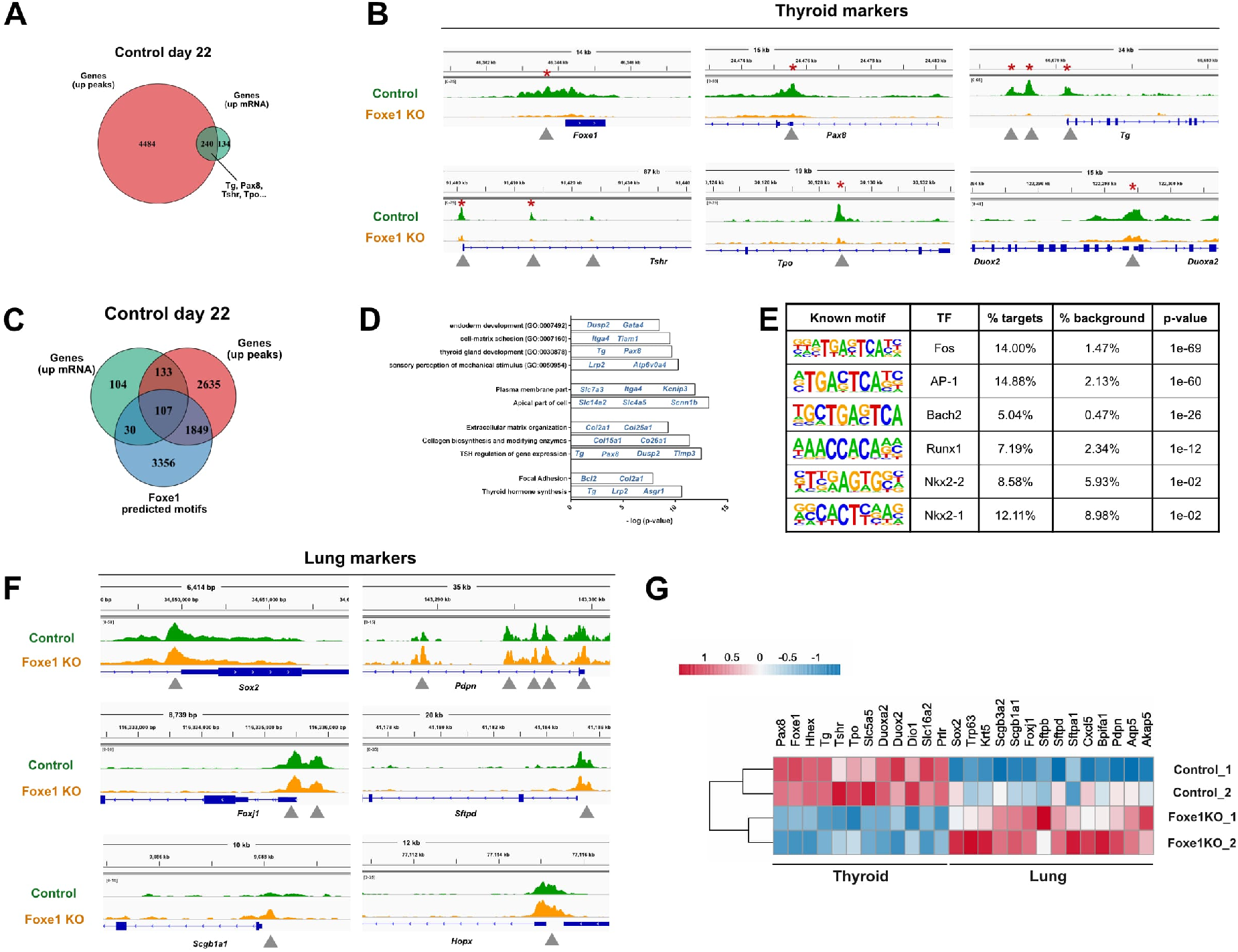
ATAC-seq profiling of Nkx2-1 expressing cells in control and Foxe1KO at day 22. (**A**) Venn diagram between the genes in which the chromatin is significantly more open and the transcripts are up-regulated in control versus Foxe1KO at day22. (**B**) Illustration of open chromatin regions at day 22 in thyroid marker genes. (**C**) Venn diagram showing 107 genes containing Foxe1 motif also being more significantly open in ATACseq and with transcripts up-regulated in control cells at day 22. (**D**) Gene Ontology (GO) and pathway functional analysis performed with 107 genes shown in D. Biological Process GO, Jensen compartments, Bioplanet and KEGG significant terms are shown in the image. (**E**) Known motifs enriched in 793 peaks more open in Foxe1KO versus control at day22; log2foldchange ≥0.58, FDR≤0.05. (**F**) Illustration of open chromatin regions at day 22 in lung marker genes. (**G**) Heatmap of log-2 transformed normalized counts from bulk RNA-seq of sorted Nkx2-1+ cells at day 22. Relevant markers for thyroid and lung lineage are shown in control and Foxe1KO samples.

To reveal putative chromatin regions regulated by Foxe1, we applied a HOMER custom motif search based on the human Foxe1 binding motif, from the Jaspar database, on total peaks of day 22 control ATAC-seq datasets (Figure 6C-D). We identified 107 genes, containing predicted Foxe1 binding motif, which are highly transcribed and located in the neighborhood of significantly up-regulated ATAC-seq peaks in the control condition (Figure 6C; Table S1). These included genes related to thyroid hormone synthesis and Tsh regulation, such as *Tg*, *Pax8*, *Dusp2*, *Timp3* and *Asgr1*, as well as various molecular players involved in endoderm development, thyroid cell polarization, and follicle organization (Figure 6D). Overall, our ATAC-seq results once again demonstrate the essential role of Foxe1 during thyroid morphogenesis and function and, additionally, provided the identification of a number of potential Foxe1 targets.

### Cis-regulatory regions around lung-related genes are equally accessible in control and Foxe1KO Nkx2-1+ cells

To investigate the gene regulatory networks enriched in Foxe1KO cells that may drive lung fate, we performed motif enrichment analysis for peaks enriched in Foxe1KO compared to control samples at day 22. We identified 793 peaks that were upregulated in Foxe1KO compared with control samples. Analysis of the HOMER - motif search revealed enrichment with Fos, AP-1 and Bach2 motifs, factors involved in apoptotic cell death, hypoxia and oxidative stress (Figure 6E) (Machado et al., 2021; Shaulian and Karin, 2002; Zhou et al., 2016). These results suggest that Foxe1 may have a broad survival function for thyroid cells in addition to inducing thyroid maturation genes.

A notable finding, however, was that none of the Foxe1KO upregulated peaks found on ATACseq analysis were directly associated with classical markers of differentiated lung cells. Indeed, the IGV plots in Figure 6F show that many cis-regulatory regions of various lung markers are equally open between control and Foxe1KO cells. Although we did not detect differences in chromatin accessibility for these genes, RNA-seq analyses of sorted Nkx2-1 cells show that they are upregulated at the level of mRNA expression in Foxe1KO, corroborating previous findings on lung formation in Foxe1 depleted cells (Figure 6G).

## Discussion

In the present study, we use our stem cell-based thyroid organoid model to show that Foxe1 is required for the proper formation of thyroid follicular structures *in vitro* and critically affects the number of thyroid follicular cells differentiated from mESCs. Furthermore, in normal thyrocytes, we identified upregulated open chromatin regions containing Foxe1 binding motifs that might be key players driving Foxe1 function. In addition to affecting thyroid differentiation, we show that a group of mESCs in a Foxe1KO context is able to maintain Nkx2-1 expression and deviate to other lineage possibilities, differentiating into lung cell types, in vitro.

### Foxe1 role in thyroid generation *in vitro*

Early thyroid development depends on the correct expression of the four major transcription factors, Nkx2-1, Pax8, Hhex and Foxe1, in the thyroid primordium. Their expression is responsible for the activation of the gene regulatory network leading to thyroid differentiation and maintenance of the differentiation status in adulthood (De Felice and Di Lauro, 2004; López-Márquez et al., 2021). Regarding Foxe1, Di lauro and coworkers have shown that although thyroid anlage is specified in the absence of Foxe1, Foxe1-null embryos exhibit severe defects in thyroid morphogenesis as early as E9.5, leading to complete disappearance of thyroid tissue around E11.5 or persistence of a small, misplaced gland (De Felice et al., 1998). In humans, homozygous mutations in Foxe1 loci lead to congenital hypothyroidism, due to severe hypoplasia or complete agenesis of the thyroid gland (Clifton-Bligh et al., 1998). Our results in the present work support the intrinsic role of Foxe1 in early thyroid development, because in the absence of Foxe1, the number of polarized thyroid follicles formed *in vitro* is drastically impaired. Moreover, the few thyroid follicular cells that differentiate in the absence of Foxe1 are abnormal and lack signs of proper thyroid maturation, as evidenced, for example, by the lack of *Nis* expression and Tg iodination. Foxe1 is thought to act downstream of Nkx2-1, Pax8 and Hhex during thyroid development, as the onset of Foxe1 expression occurs through the direct action of Pax8, suggesting a hierarchy among thyroid transcription factors (López-Márquez et al., 2021; Parlato et al., 2004). Indeed, we have shown that transient induction of mESCs with Nkx2-1/Pax8 is sufficient to induce Foxe1 expression *in vitro* (Antonica et al., 2012). The fact that Foxe1 is downstream in the gene regulatory network for thyroid specification might explain why follicle-like cells are still present in the absence of Foxe1, even if they are scarce and non-functional.

One of the proposed mechanisms of Foxe1 action on thyroid morphogenesis is the direct regulation of genes critical for thyroid maturation and function. In differentiated thyroid follicular cells, Foxe1 is involved in TSH-mediated regulation of *Tg* and *Tpo* expression and Foxe1 binding sites have been described in the promoters of these genes (Aza-Blanc et al., 1993; Francis-Lang et al., 1992; López-Márquez et al., 2019; Santisteban et al., 1992). Foxe1 can also bind directly to cis-regulatory regions of Nis/Slc5a5 and Duox2 (Fernández et al., 2013). In the present work, we show that the expression of many thyroid functional genes is impaired in the absence of Foxe1, confirming these previous reports. No cells expressing *Nis*, *Tpo* and *Duox2* were identified by scRNAseq analyses of Foxe1KO cells. Integrated bulk ATAC/RNA-seq analyses showed that reduction in expression of genes involved in thyroid maturation might be associated with silencing of many cis-regulatory regions, including at their promoters. Interestingly, there are 17-fold more opened chromatin regions in Nkx2-1+ control cells compared with Foxe1KO/Nkx2-1+ cells, indirectly supporting the view that Foxe1 has an impact on chromatin remodeling in thyrocytes. Although our experiments were not designed to explore this aspect, the function of Foxe1 as a pioneer factor, well described for other members of the Forkhead family of transcription factors (Zaret and Carroll, 2011), has previously been suggested in relation to its ability to bind the compacted chromatin around the inactive *Tpo* promoter (Cuesta et al., 2007).

In addition, we identified novel genomic targets of Foxe1 by direct comparison analyses between Foxe1-depleted and control Nkx2-1-expressing cells. *In silico* prediction of Foxe1 binding motifs in up-regulated genes at the level of RNA expression and chromatin accessibility revealed 107 potential targets of Foxe1, including *Pax8* and *Tg* (Francis-Lang et al., 1992). Other targets identified included genes involved in proper thyroid function and TSH receptor activity or deregulated in thyroid cancer, such as *Lrp2/Megalin* (Marinò and McCluskey, 2000), *DREAM/Kcnip3* (Andrea et al., 2005), *Timp3* (Zarkesh et al., 2018), *Slc26a7* (Dom et al., 2021) and *Bcl-2* (Fagman et al., 2011; Porreca et al., 2012). Moreover, Foxe1 binding motifs were also found in genes related to the maintenance of follicle structure, the expression of various membrane-bound transporters, and the regulation of cell-matrix adhesion. In summary, the present study sheds light on novel molecular players essential for thyroid development and homeostasis. Ultimately, our thyroid stem cell-derived organoid system proved to be a powerful model for such analyses because 1, it is an *in vitro* model, so obtaining an adequate amount of cells is not as limited as *in vivo*; 2, growing differentiated thyrocytes in 3D, to generate polarized follicular structures, allows the correct localization and signaling of factors involved in the thyroid hormone synthesis machinery, and therefore the screening of new modulators of thyroid function may be more representative.

### Lung generation *in vitro*, in the absence of Foxe1

Besides the disruption of thyroid lineage formation in Foxe1KO cells, we additionally observed the formation of Nkx2-1+ organoids that lack Pax8 expression and are morphologically distinct from our stem cell-derived thyroid follicles. Considering that Nkx2-1, in endoderm-derived tissues, is also present in embryonic and adult lung cells (Herriges and Morrisey, 2014; Lazzaro et al., 1991; Nikolić et al., 2018), we hypothesize that permissive signals for lung differentiation occurred in the context of Foxe1 depletion. Indeed, these Nkx2-1+ organoids possess numerous markers of airway cell types. Basal lung cells, for example, could be identified in Krt5+/Sox2+/Trp63+/Nkx2-1+ cystic organoids. Other airway-like organoids are organized as branched structures and consist of a Sox2+ single-layered epithelium containing goblet-like cells that secrete mucus in the luminal space. Finally, characterization of lung organoids by scRNAseq revealed the presence of alveolar-like cells, which confirmed our ultrastructural microscopic analyses. Overall, in the absence of Foxe1 and using the same protocol to induce thyroid lineage, a large subset of mESCs can diverge to a lung fate and differentiate into a mixture of alveolar and airway-derived cell types, with the latter being more prevalent.

Nkx2-1 is the earliest known marker of respiratory fate. As thyroid, future lung progenitors are derived from the ventral anterior foregut endoderm, posterior to the thyroid field, and the onset of lung differentiation can be detected by the expression of Nkx2-1 (Cardoso and Lu, 2006; Lazzaro et al., 1991). We speculate that in our current model system, since we directly overexpress Nkx2-1 (and Pax8) without stepwise restriction of lineage commitment, a subset of mESCs are more prone to activate a lung differentiation program that is quiescent under control conditions, but the absence of a key gene for thyroid differentiation leads to a lineage shift in these cells. Our study supports the hypothesis of a certain level of fate plasticity of early Nkx2-1 progenitors generated *in vitro*. Interestingly, an elegant study has recently demonstrated that a subset of NKX2-1+ progenitors, derived from human pluripotent stem cells, can deviate to alternative, non-lung endodermal cell fates (Hurley et al., 2020).

It is important to point out that our bulk ATAC-seq analyses suggest that at the level of chromatin accessibility, the main differences between Foxe1KO and control Nkx2-1+ cells are mainly due to an enrichment of a thyroid differentiation program towards the end of the differentiation protocol: cis-regulatory regions of genes such as *Tg*, *Tshr*, *Tpo*, *Duox2*, *Duoxa2* are significantly more open in control versus Foxe1KO- Nkx2-1+ cells. On the other hand, the chromatin state around genomic locations of lung-typical genes, including promoter regions, appears to be equally open in both conditions, even if these genes are more highly expressed in Foxe1KO Nkx2-1+ cells. Two hypotheses can be inferred from these results: 1, Many known lung markers, such as *Sftpb*, *Scgb1a1*, *Napsa*, *Ager*, *Aqp5*, and *Sox9*, are also expressed to some extent in thyroid, so it makes sense that chromatin accessibility does not differ substantially between these cells for these genes (Dame et al., 2017; Silberschmidt et al., 2011; and our scRNAseq). Further differential analyses throughout mouse development are needed to better understand the role of these genes in a thyroid context (Ikonomou et al., 2020; Mou et al., 2012). 2, By default, mESC-derived Nkx2-1 cells may have some potential to give rise to lung cells. Depletion in Foxe1 expression would allow this potential to be unleashed. It is interesting to see that, in our results, the global chromatin accessibility of in vitro Nkx2-1 progenitor cells is very similar at early stages (day 10), suggesting a direct role of transient overexpression of Nkx2-1 and Pax8 in triggering both thyroid/lung programs. Such an event of concomitant generation of two endodermal lineages has been demonstrated earlier in the direct lineage conversion of fibroblasts to liver and large intestine cells, following overexpression of Foxa1 (Morris et al., 2014).

Finally, we must emphasize that although Pax8 is also forcibly expressed by doxycycline treatment, we do not assume that Pax8 directly mediates the induction of lung fate in our model, which is supported by *in vivo* evidence showing that Pax8 is not expressed in the lung during mouse embryogenesis (Kurmann et al., 2015; Mou et al., 2012). Pax8 is rapidly downregulated in Foxe1KO cells, while Nkx2-1 expression, albeit less pronounced than under control conditions, is maintained throughout the protocol. Moreover, no derivation of thyroid or lung is achieved in Pax8KO mESC lines (data not shown). In other words, simple removal of another key transcription factor for thyroid differentiation did not result in the appearance of a lung instead of a thyroid, as is the case in Foxe1. Nevertheless, we speculate that in the case of Pax8 loss of function in mESCs, activation of high endogenous Nkx2-1 levels is less successful, suggesting that Pax8, in our model (Antonica et al., 2012), helps to increase Nkx2-1 expression in the first days, and thus Pax8 might be indirectly involved in the initiation of a lung program *in vitro*.

In conclusion, the present work improves our understanding of the crucial role of Foxe1 in triggering correct thyroid tissue formation and function, and points to novel molecular players for further research. On the other hand, this report also evidences the intricate relationships among endodermal lineages, as it seems to be the case for thyroid and lung, especially during differentiation. Whether transient thyroid/lung bipotent progenitors do exist *in vivo*, in a similar phenomenon observed during liver and pancreas lineage derivation for example (Deutsch et al., 2001; Xu et al., 2011), would be an interesting avenue for further investigations. We speculate that Foxe1 could be one of the factors orchestrating this lineage choice.

## Limitations of the study

From our knowledge, the present manuscript is the first to demonstrate lung lineage derivation at the expense of thyroid lineage, due to the absence of a transcription factor. However, additional work is required to fully explore those interesting findings. Although our ATACseq analysis aided to unveil some aspects towards thyroid biology, the same is not possible to conclude regarding lung generation in the absence of Foxe1. The heterogeneity of Foxe1KO-derived cells, associated with the low resolution of bulk ATAC-seq transcriptomics, may have hampered the scrutiny of this aspect. The use of new tools such as scATACseq may be necessary in the future. Moreover, the investigation of the developmental competence of early progenitors expressing Nkx2-1, in the gut tube, regarding their chromatin state (Wang et al., 2015), is an interesting avenue for future research. Early Nkx2-1 cells in the thyroid anlage are still difficult to obtain *in vivo* (Ikonomou et al., 2020), but future technologies may aid in this aspect. Finally, the present study was made entirely *in vitro*, and as such, contains the limitations inherent of this strategy.

## Acknowledgements

We acknowledge the ULB flow cytometry platform (Christine Dubois), the ULB genomic core facility (F. Libert and A. Lefort), LiMIF platform for confocal microscopy (J.-M. Vanderwinden) and Veronique Janssens for lab management and technical help. M. Saiselet for help with 10X genomics assay, D. Frank (University of Pennsylvania) for providing scRNAseq metadata of mouse lung samples, H. Lasolle for RNA-seq discussions, Y.Song for help in ATACseq analysis. Work in S.P.S.’s lab is supported by MISU (34772792) and MISU-PROL (40005588) funding from the FNRS and from Fondation Jaumotte-Demoulin. This work was supported in part by grant DK15070 from The National Institutes of Health. USA, to SR.

## Author contributions

B.F., C.B., S.C designed the experiments and analyzed the data. M.R., S.P.S and S.E.E. performed additional data analysis. B.F., C.B., M.R. generated genetic tools. C.B., M.R. performed *in vitro* organification and C.B., P.G. confocal microscopy. B.F., S.P.S, S.E.E. analyzed scRNAseq data. B.F., S.P.S performed ATACseq analysis.

D.M. performed electron microscopy, D.P.M analyzed EM data, S.C. and S.R. acquired funding for the project. B.F., S.C. wrote the manuscript and all authors read, edited and approved the final manuscript.

## STAR Methods

### KEY RESOURCES TABLE RESOURCE AVAILABILITY

#### Lead contact

Further information and requests for resources and reagents should be directed to and will be fulfilled by the lead contact, Sabine Costagliola

#### Materials availability

The mESC lines and plasmids generated in the current study are available upon request.

#### Data and code availability

Original data associated with this study is deposited in the NCBI Gene Expression Omnibus under accession numbers GSE182337, GSE182480 and GSE182676. Information regarding published datasets used in the present paper can be found in Franck et al, 2019. Source data are provided with this manuscript. scRNAseq data and gene expression profile (interactive tool) can be accessed at https://barbaraffonseca.shinyapps.io/Foxe1KO/.

### METHOD DETAILS

#### mESCs culture and differentiation

All mESC lines used in the present paper are derived from the A2Lox.Cre_TRE-Nkx2-1/Pax8_Tg-EGFP mESCs line previously generated by our group (Romitti et al., 2021). Maintenance and differentiation of mESCs lines were performed as previously described (Antonica et al., 2012, 2017). Briefly, mESCs were cultured on a feeder-layer of irradiated mouse embryonic fibroblasts, in maintenance medium. For the differentiation protocol, mESCs were isolated from feeder-layer cells, counted and cultured in hanging drops (1,000 cells per drop) for generation of embryoid bodies (day 0). Four days later, embryoid bodies were collected, embedded in growth factor-reduced Matrigel and 50µl Matrigel drops re-plated into 12-well plates. Embryoid bodies were differentiated and cultured using the differentiation medium supplemented with 1 mg/ml Doxycycline, for induction of exogenous Nkx2-1 and Pax8 transgenes for three days (day4-day7), followed by two weeks treatment (day7-day22) with 1mU/ml hrTSH or 300µM 8-br-cAMP, as indicated.

#### Generation of mESC lines by TALEN technology

TALEN technology was used to edit the A2Lox.Cre_TRE-Nkx2-1/Pax8_Tg-EGFP mESC line to generate FoxE1KO cells. TALENs synthesis was performed by using Golden Gate Technology (Weber et al., 2011). mESCs were seeded into 10 cm petri dishes (150.000 cells/petri) and, in the next day, transfected with 7 µg of each TALEN-encoding plasmid (designed to target the Forkhead domain) and 7µg of a fluorescent surrogate reporter plasmid allowing enrichment of cells with nuclease induced-mutations (GFP reporter Assay) (Ma et al., 2013). Cell transfection was performed with Lipofectamine3000 according to the manufacturer’s instruction. 48h later, cells were trypsinized and resuspended in PBS containing 2% of embryonic-stem-certified fetal bovine serum for cell sorting (FACS Aria and FACSDiva Software). GFP+ cells were seeded in 96-well plates in order to have 1 GFP+ cell per well. Individual colonies were expanded in 24-well, 12- well or 6-well plates with maintenance medium during one week. A second cell sorting procedure was performed in individual clones to obtain a fully pure population of mESCs, devoid of feeder-layer cells, for genomic PCR analysis. Sequences of TALEN constructs are depicted in Table S2.

#### Generation of Nkx2-1 endogenous reporter mESC line by CRISPR/Cas9

For production of single-guide RNAs (sgRNAs) targeting the 3’-UTR region of Nkx2-1 loci, a fragment of the desired genomic mouse sequence was obtained from NCBI and used as input in the CRISPOR webtool (http://crispor.tefor.net/ (Concordet and Haeussler, 2018)). The most suitable sgRNA was chosen and checked for possible off-targets (Table S2). Forward and reverse oligonucleotides, corresponding to the sgRNA sequence, were designed for insertion into the pU-BbsI-T2A-Cas9-BFP plasmid (Chu et al., 2015). To generate the pUC57-mNkx2-1-T2A-mKO2-pGK-Puro targeting vector, approximately 1000bp of each left and right Nkx2-1 homology arms, around predicted sgRNA cutting-site, were PCR amplified and inserted in a template plasmid, containing the T2A-mKO2 and loxP-flanked pGK-Puro selection cassette. Lipofectamine3000 was used to transfect mESCs, according to the manufacturer’s instruction. 48h later, cells were sorted based on BFP+ expression and seeded at the clonal level in 96-well plates, as described above. Positive selection of puromycin-resistant clones was performed by treatment with 1µg/ml puromycin for 72 hours. Integration of desired cassette was confirmed by PCR of genomic DNA. Successful targeting was achieved in 7 among 125 screened clones. The selected clones integrated the donor template in both alleles. Excision of loxP-flanked pGK-Puro fragment was achieved later by transfection with pSalk-Cre plasmid, followed by a series of cell cloning, expansion and negative selection based on susceptibility to puromycin treatment. Genomic removal of PuroR sequence was confirmed by PCR. Finally, mKO2 reporter activation and compatibility with Nkx2-1 protein expression was verified. For clarity, in the present paper, the novel line generated (i.e. A2Lox.Cre_TRE-Nkx2-1/Pax8_Tg-EGFP_Nkx2-1-T2A-mKO2 mESCs line) is named as “Nkx2-1 reporter line”.

#### Generation of Foxe1KO/Nkx2-1 reporter mESC line

SgRNAs targeting mouse Foxe1 coding sequence were chosen based on suitable sgRNAs prediction from CRISPOR website and insertion of sgRNAs sequences into the pU-BbsI-T2A-Cas9- BFP plasmid was performed as cited above. Two sgRNAs were chosen (one targeting the Forkhead domain, and the other outside) and two Foxe1KO lines, targeting distinct genomic domains of Foxe1, were produced (Table S2). Nkx2-1 reporter mESC cells were transfected, BFP sorted, clonal seeded and selected based on PCR and Sanger sequencing to detect clones containing frameshift mutations in Foxe1 loci. A second cell sorting procedure was performed in individual clones to obtain a fully pure population of mESCs, devoid of feeder-layer cells, for genomic PCR analysis.

All cell lines here generated were validated, in at least two different clones, by maintenance of pluripotency cell markers, ability to spontaneous differentiate into the three germ layers, activation of tetracycline-inducible Nkx2-1-Pax8 transgene and reproducibility regarding in vitro differentiation. Regarding Foxe1KO/Nkx2-1 reporter lines, similar results were obtained in Foxe1KO lines derived from distinct designed. Only one is shown in the present paper.

#### PCR detection of mutated clones (TALEN/CRISPR-Cas9)

Genomic DNA was extracted from individual mESCs clones, using a genomic DNA lysis buffer and procedure as described (Verma et al., 2017). PCR was performed using Q5 High-Fidelity Taq Polymerase according to the manufacturer’s instructions. 100 ng of genomic DNA samples were used in all reactions. Primer sets were used to amplify targeting and cutting site of designed TALENs/CRISPR-Cas9 guide RNAs. PCR products were then directly cloned in a TOPO-Blunt plasmid by using Zero Blunt® PCR Cloning Kit.

#### RNA extraction and RT–qPCR

For RNA preparation, cells were lysed in RNeasy Lysis buffer + 1% β-mercaptoethanol, and total RNA was isolated using RNeasy microRNA preparation kit according to the manufacturer’s instructions. Reverse transcription was done using Superscript II kit. RT-qPCR was performed in technical triplicates using KAPA SYBR Fast qPCR kit and a CFX Connect Real-Time PCR System. Results are presented as linearized values normalized to the housekeeping gene β2m-Globulin and the indicated reference value (2-ΔΔCt). Moreover, relative expression of each target gene is presented as fold-change compared to untreated cells (-dox condition). The gene-expression profile was confirmed in at least two different clones of each cell line. Primers used are listed in Table S3.

#### Scanning Electron microscopy

Organoids embedded in Matrigel were fixed in glutaraldehyde 2.5% overnight at 4°C, rinsed, and embedded in agarose 4%. Sections of 300µm were produced in a Vibratome (Leica) and post-fixed in OsO4 (2%) for 1 hour. All treatments were done in 0.1M cacodylate buffer (pH 7.2). After serial dehydration in ethanol, samples were dried at critical point and coated with platinum by standard procedures. Observations were done in a Tecnai FEG ESEM QUANTA 200 at 30kV and images were acquired and processed by SIS iTEM software.

#### Iodide organification assay

The iodide organification assay was performed as previously optimized for mouse thyroid organoids (Antonica et al., 2012). Briefly, samples were rinsed with HBSS and incubated with 1ml per well of organification medium (1,000,000 c.p.m. per ml of ^125^I and 100nM sodium iodide in HBSS for 2h at 37°C. Next, 1ml of a TPO inhibitor (4mM methimazole) was added and cells were dissociated with 0.1% Trypsin/1mM EDTA for 15min. Radioactivity, derived from iodide uptake by thyrocytes, was measured with a γ-counter. To quantity the radioactivity of protein-bound ^125^I (PBI), proteins were precipitated in a solution containing 1mg γ-globulins and 2ml of 20% TCA followed by centrifugation at 2000r.p.m. for 10min. Iodide organification was calculated as iodide uptake/PBI ratio.

#### Immunofluorescence and immunohistochemistry

Cells were fixed in 4% formaldehyde solution for 30min and washed three times five minutes in PBS. Then, cells were blocked in PBS containing 3% bovine serum albumin (BSA), 5% horse serum and 0.3% Triton X-100 for 30min at room temperature. The primary and secondary antibodies were diluted in PBS containing 3% BSA, 1% horse serum and 0.1% Triton X-100. Primary antibodies were incubated overnight at 4°C followed by incubation with secondary antibodies for 1h30 at room temperature. Nuclei were stained with 4′,6-diamidino-2-phenylindole. Coverslips were mounted with Glycergel.

#### Flow cytometry intracellular immunostaining

Nkx2-1 reporter and unmodified mESCs were treated up day 14 of thyroid differentiation protocol and prepared for flow cytometry immunostaining as follows: Matrigel-drops were first digested with a HBSS solution containing 10 U/ml dispase II and 125 U/ml of collagenase type 1A for 30 minutes at 37°C; then, cell dissociation was performed with TripLE Express for maximum 15min, to obtain a single cell suspension. After centrifugation, samples were PBS rinsed and fixed in 1.6% formaldehyde solution in PBS for 15min at room temperature, followed by a step of permeabilization with 0.1% Triton in PBS for 1min at 4°C. Before staining, permeabilized samples were blocked for 10 min in a solution containing 4% horse serum and 0.5% Tween 20. Primary anti-rabbit Nkx2-1 antibody was diluted at 1:100 ratio in a 0.5% Tween washing solution and added to samples for 30 min at 4°C. They were then rinsed three times with washing solution (0.5% Tween in PBS) before incubation for 30min with Cy5-conjugated anti-rabbit antibody diluted at 1:300 ratio in 0.5% Tween in PBS. Additional washes were performed with the washing solution before addition of a live cell marker (Calcein Violet 1µM) and data acquisition using a LSRFortessa X-20 flow cytometer and FACSDiva software. Flow cytometry controls such as unstained cells, isotype controls and negative controls (undifferentiated cells: “-dox control”) were included in all experiments.

#### RNA isolation and RNA-seq analysis

For preparation of bulk RNA-seq samples, Foxe1KO and control cells from Nkx2-1 reporter line were cultured following the differentiation protocol and cell suspension was obtained as above described (“Flow cytometry intracellular immunostaining” section). Nkx2-1+ (mKO2+) cells were sorted (FACS Aria; BD Bioscience) at day10 and day22. 10000 mKO2+ cells per condition were collected directly into 700µl of Qiazol lysis reagent and RNA isolation performed with miRNeasy micro kit following manufacturer’s indications. The quality and quantity of the resulting RNA was then tested using Bioanalyser 2100 (Agilent) and RNA 6000 Nano Kit. RNA integrity was preserved (RIN=8.5) and no genomic DNA contamination was detected. Ovarion Solo RNA-seq Systems was employed, as indicated by the manufacturer, to produce high quality indexed cDNA libraries, which were quantified using Quant-iT PicoGreen kit and Infinite F200 Pro plate reader (Tecan); DNA fragment size distribution was examined on 2100 Bioanalyser (Agilent) using DNA 1000 kit. Normalized and pooled indexed libraries (10ρM) were loaded on flow cells and sequenced on the HiSeq 1500 system (Illumina) in a high output mode using HiSeq Cluster kit v4. Approximately 10 million of 125nt-long paired-end reads were obtained for each library. After removal of low-quality bases and Illumina adapter sequences using Trimmomatic software (Bolger et al., 2014), sequence reads were aligned against mouse reference genome (Grcm38/mm10) using Hisat2 software with default parameters (Kim et al., 2015). Raw counts were obtained using HTSeq software (Anders et al., 2015) using Ensembl genome annotation GRCm38.87. Normalization, differential expression and Gene Ontology analyses were performed with at least two biological replicates per sample, using website iDEP version 0.92 (Ge et al., 2018). Additional Gene Ontology analysis and identification of statistically significant terms (p < 0.05) was performed with Enrichr (Kuleshov et al., 2016).

#### Single cell RNA-seq preparation and sequencing

At day22 of the differentiation protocol, cell populations derived from Foxe1KO/Nkx2-1 reporter mESC lines were isolated for scRNAseq profiling. Culturing and preparation of cell suspension for FACS-sorting were performed as mentioned above for bulk RNA-seq. Different proportions of EGFP+, mKO2+ and mKO2- cells were sorted to guarantee representation of various cell types in the profiled sample (15%, 50%, 35%, respectively). Sorted cells were collected in PBS at a density of 800cells/ul and diluted accordingly to kit’s instruction (10x Genomics Chromium Single Cell 3’ v3). 12000 cells were loaded onto a channel of the Chromium Single Cell 3′ microfluidic chip and barcoded with a 10X Chromium controller. Subsequently, RNA was reverse transcribed and amplified according to manufacturer’s recommendations. Library preparation (e.g. fragmentation, dA tailing, adapter ligation, indexing PCR) was performed based on 10x Genomics guidelines. Libraries were sequenced on a Illumina NovaSeq 6000 system.

#### Single cell transcriptomic data analysis

Raw sequencing data was aligned and annotated against the Grcm38/mm10 mouse reference genome, in which mKO2 and EGFP sequences were added. Cell Ranger Software (v.2.1.0), provided by 10x Genomics, was used for demultiplexing with default parameters. The raw counts generated from 10x Chromium pipeline were clustered using R Seurat package (version 3.2.0) (Stuart et al., 2019). Briefly, quality control pre-processing was performed to keep cells passing the following criteria: had between 1500 and 58000 UMI counts, showed expression of at least 750 unique genes and had less than 10% of UMI counts corresponding to mitochondrial genes. The remaining data was log-normalized, regressed out to remove effects of library size and enrichment of mitochondrial and cell cycle related genes, and scaled, using *SCTransform* function. Principal component analysis (PCA) was calculated using the expression data of the most variable genes and the first 10 principal components were used to graph-based clustering and UMAP plot visualization. Different values in the resolution variable (*FindClusters* function) were tested and resolution of 0.7 was used for clustering. Differentially expressed genes were computed with the *FindAllMarkers* function and used for heatmap visualizations. To better identify specifically lung cells, cells expressing *Nkx2-1* and *Epcam*, but devoid of *Thyroglobulin* and *Pax8* expression, were extracted from original dataset, re-clustered based on variable genes and new UMAP visualization obtained. Signature scores based on expression of selected genes were calculated using *AddModuleScore* function. Gene lists can be found on Table S1.

#### Integrative single-cell RNAseq analysis

For better understanding of Foxe1KO effect, we compared Nkx2-1+ cells derived from Foxe1KO line with a previous scRNAseq dataset performed in a control mESC line (Romitti et al., 2021). For this, we extracted Nkx2-1+ cells from both datasets, combined in an unique object and calculated pairwise correspondences between individual cells using Integration features from R toolkit (Stuart et al., 2019). After integration, downstream analyses such as graph-based clustering and UMAP dimensionality reduction were performed as described above. In addition, to identify shared cell populations among in vivo mouse lung and Foxe1KO-derived lung organoids, *Nkx2-1*+/*Epcam*+/*Tg*-/*Pax8*- cells from Foxe1KO dataset were compared to E17.5 mouse lung (Frank et al., 2019). For this, Seurat object of E17.5 mouse lung scRNAseq dataset was updated and subjected to the same normalization and scaling process described above (*SCTransform* function). For better visualization purposes, the original dataset was downsampled to 2000 cells before integration.

#### ATAC sequencing

Biological replicates were obtained from control and Foxe1KO cells (Nkx2-1 reporter line), at different points of our differentiation protocol. 50000 Nkx2-1(mKO2+) cells were sorted from both lines at day 10 and immediately proceed to sample preparation, based on Omni-ATAC protocol (Corces et al., 2017). Cells derived from embryoid bodies before (day 4) and after doxycycline treatment (day7) were also collected to distinguish open chromatin regions related with tetracycline-induced exogenous transgene activation. After centrifugation, cell pellets were resuspended in 50µl of an ice-cold cell lysis buffer (0.1% Igepal, 0.1% Tween20 and 0.01% Digitonin in Omni-ATAC Resuspension buffer). After 3 min, samples were centrifuged for 15min at 800g and subsequently, nuclei were resuspended in 50µl of reaction buffer (2.5μl Tn5 transposase, 22.5μl TD buffer, both from Nextera DNA sample preparation kit; 16.5μl PBS, 0.5μl 1%Digitonin, 0.5μl 10% Tween20 and 5μl H20). Tagmentation reaction was performed for 30min at 37°C in a rocking plate (1000 rpm). DNA was purified using the MiniElute purification kit following the manufacturer’s indications. DNA libraries were PCR amplified, DNA quality verified on 2100 Bioanalyser (Agilent) using DNA 1000 kit and size selected from 200 to 800 bp, following the manufacturer’s recommendations.

#### ATAC-seq analysis

For main steps of pre-processing and mapping of ATACseq data, a local installation of Galaxy platform was used (use.galaxu.eu; (Afgan et al., 2018)). Briefly, adapter sequences were removed with Trim Galore, using default parameters. ATACseq paired-end reads were aligned to mouse genome Grcm38/mm10 with Bowtie 2, modifying default parameters to include fragments of up 1000bp, allowing dovetailing and using “very sensitive option” preset. Subsequently, mitochondrial genes, bad quality mapped sequences and PCR duplicates were removed. Peak calling was performed for each sample using MACS2, with parameters setting of -q 0.05 and -- shift 0. Peaks from all samples were merged for downstream analysis. Scaling of bam files generated by MACS2 were performed and used for visualization of data tracks with Integrative Genomics Viewer (IGV). Moreover, files derived from MACS2 peak calling were used as input for Differential Binding Analysis using DiffBind package in R (Stark and Brown, 2011). Annotation of nearest genes associated with differentially regulated genomic regions was performed using HOMER (Heinz et al., 2010). Two biological replicates were used for sample and significant differential peaks are were filtered according to these criteria: log2 fold change ≥0.58 and false discovery rate (FDR) ≤0.05. De novo motif search was performed using findMotifs.pl from HOMER package with default parameters. To obtain Foxe1 enriched motifs in total ATACseq peaks, the human Foxe1 motif (MA1487.1) obtained on Jasper database (Castro-Mondragon et al., 2021), was used as a query in findMotifs.pl command.

#### Statistical analysis

For most experiments (RT-qPCR, iodide organification, immunofluorescence and flow cytometry), at least two different wells per condition were used in each differentiation experiment. Furthermore, at least three independent experiments were performed. Statistical significance was tested as follows: two-group comparison by unpaired t-test and multiple-group comparison by the one-way analysis of variance test with a post-hoc Tukey’s comparison test. *P<0,05, **P<0,01, ***P<0,001. Bar plots show mean ± s.e.m., unless otherwise indicated. GraphPad Prism version 6 was used for most analyses. Shapiro-Wilk normality test and Mann-Whitney unpaired rank test were used in R to test significance of signature score enrichment in scRNAseq datasets comparisons.

#### Imaging

Fluorescence imaging was performed on a Leica DMI6000 with DFC365FX camera and a ZeissLSM510 META confocal microscope. Affinity Designer and ImageJ software (Schindelin et al., 2012) were used to adjust brightness, contrast and picture size.

## SUPPLEMENTAL INFORMATION

### Supplementary Figures titles and legends

**Figure S1.**
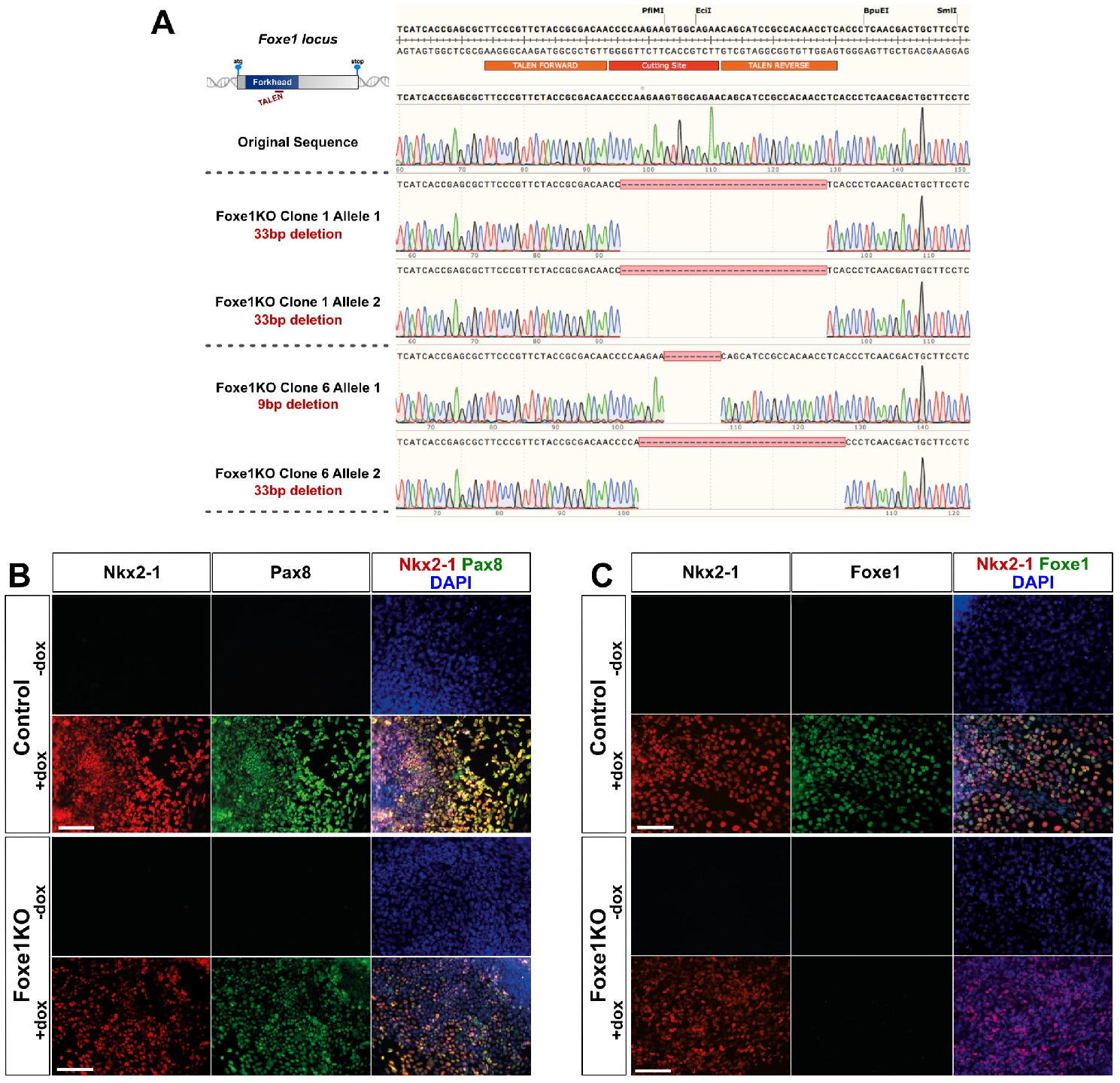
Generation and validation of Foxe1 KO mESC lines. (A) Genomic profiling of two Foxe1 KO mESC clones. One clone presents an homologous deletion of 33 bp and the second one possess an heterozygous deletion of 9 and 33 bp at Foxe1 loci, respectively. (**B-C**) Immunostaining of control and Foxe1KO mESCs after Dox-mediated induction of Nkx2.1-Pax8 during 3 days (d4-d7). (B) The Dox-inducible co-expression of NKX2.1 and PAX8 is not altered by genome editing manipulations in mutated clones. (**C**) Foxe1KO mESCs express NKX2.1 while Foxe1 protein expression is abolished. Scale bars : 150 µm.

**Figure S2.**
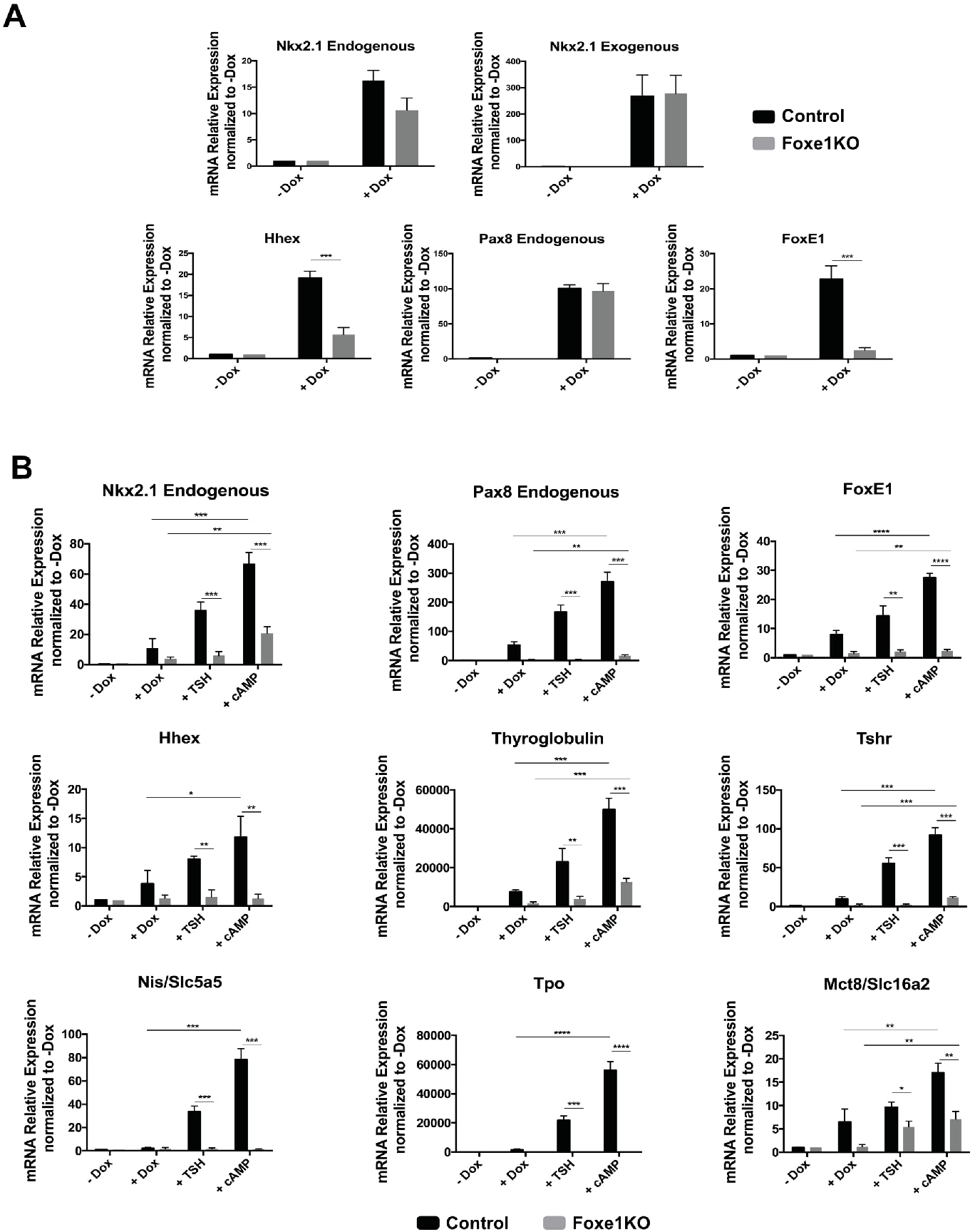
Foxe1 depletion in mESCs abolishes thyroid follicle differentiation. RT-qPCR analyses of thyroid-expressed genes in control and Foxe1KO cells at day 7 (A) and day 22 (**B**) after Dox-mediated induction (day4-day7) followed by TSH (+TSH) or 8-br-cAMP (+cAMP) treatment until day 22. (**A**) At day 7 of differentiation, downregulation of some thyroid genes such as Hhex and Foxe1 is already observed in Foxe1KO cells, whereas Nkx2-1 and Pax8 endogenous expression is not affected. Observe that Dox-mediated induction of Nkx2-1/Pax8 was successful since exogenous expression of Nkx2-1 is not affected in both lineages. (**B**) Expression of endogenous thyroid genes at day 22 are drastically reduced in Foxe1KO cells (+TSH and +cAMP conditions) in comparison with the control line. Relative expression of each transcript is presented as fold change compared to untreated cells (-Dox) as mean +/-sem. Unpaired t-test was used for statistical analysis. *P<0,05, **P<0,01, ***P<0,001. Dox: doxycycline.

**Figure S3.**
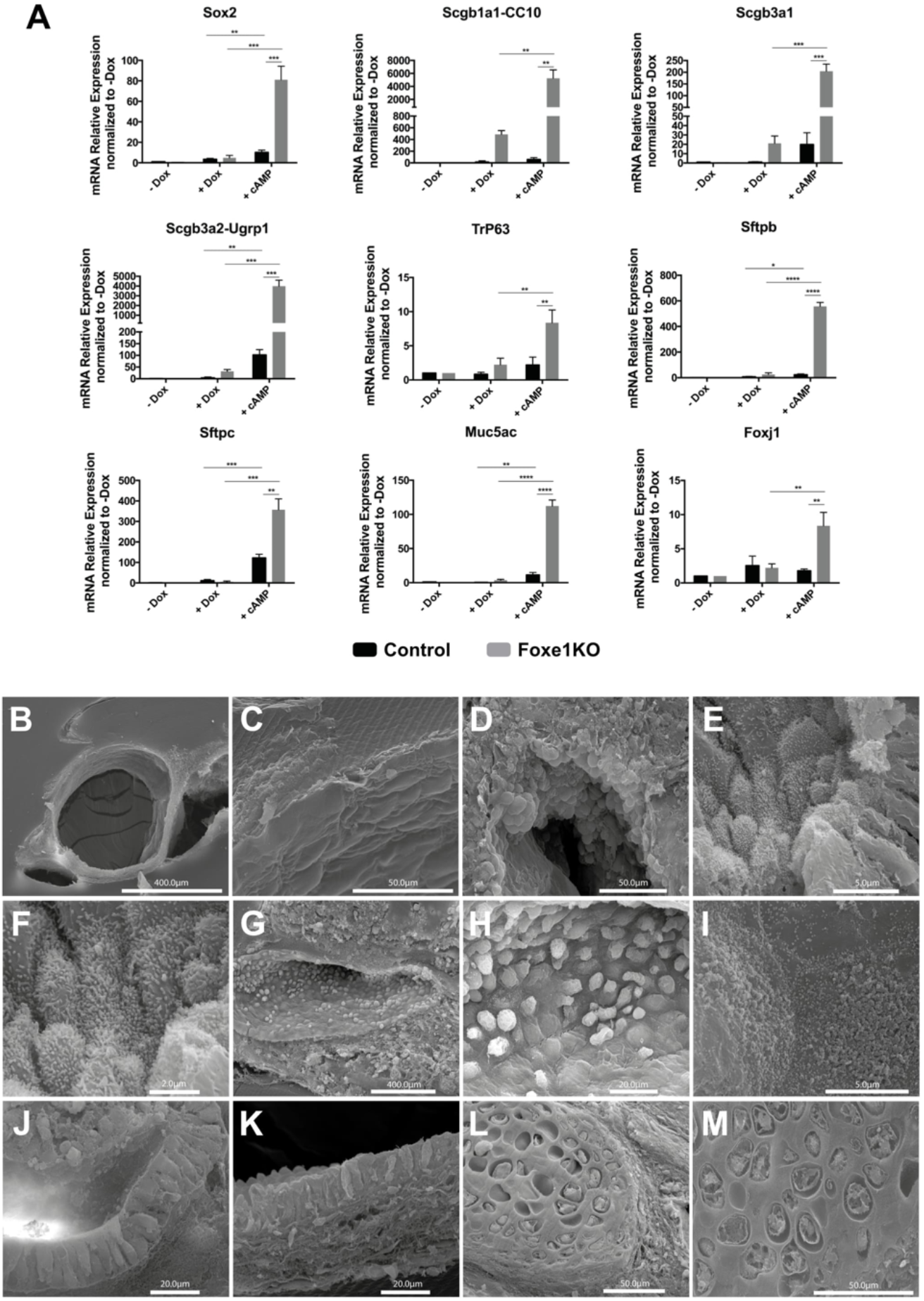
*In vitro* generation of pulmonary structures. RT-qPCR and scanning electron microscopy of *in vitro*- generated Foxe1KO pulmonary structures after Dox-mediated induction of Nkx2.1- Pax8 followed by 8-br-cAMP treatment until day22. (**A**) mRNA expression of endogenous relevant pulmonary genes at day 22. Observe up- regulation of lung-related markers in the +cAMP condition compared with untreated cells (-Dox) and dox-induced only condition (+Dox). Relative expression of each transcript is presented as fold change compared to untreated cells (-Dox) as Mean +/-SEM. Unpaired t-test was used for statistical analysis. *P<0,05, **P<0,01, ***P<0,001. (**B-M**) Scanning electron microscopy shows the presence of a large panel of pulmonary structures and differentiated cell types: branched pulmonary structures with a layer of pneumocytes type I (**C**), goblet cells secreting mucus (**D-F**), multiciliated cells (**G-I**), pyramidal pulmonary epithelium (**J-K**), chondrocytes and associated cartilage (**L-M**).

**Figure S4.**
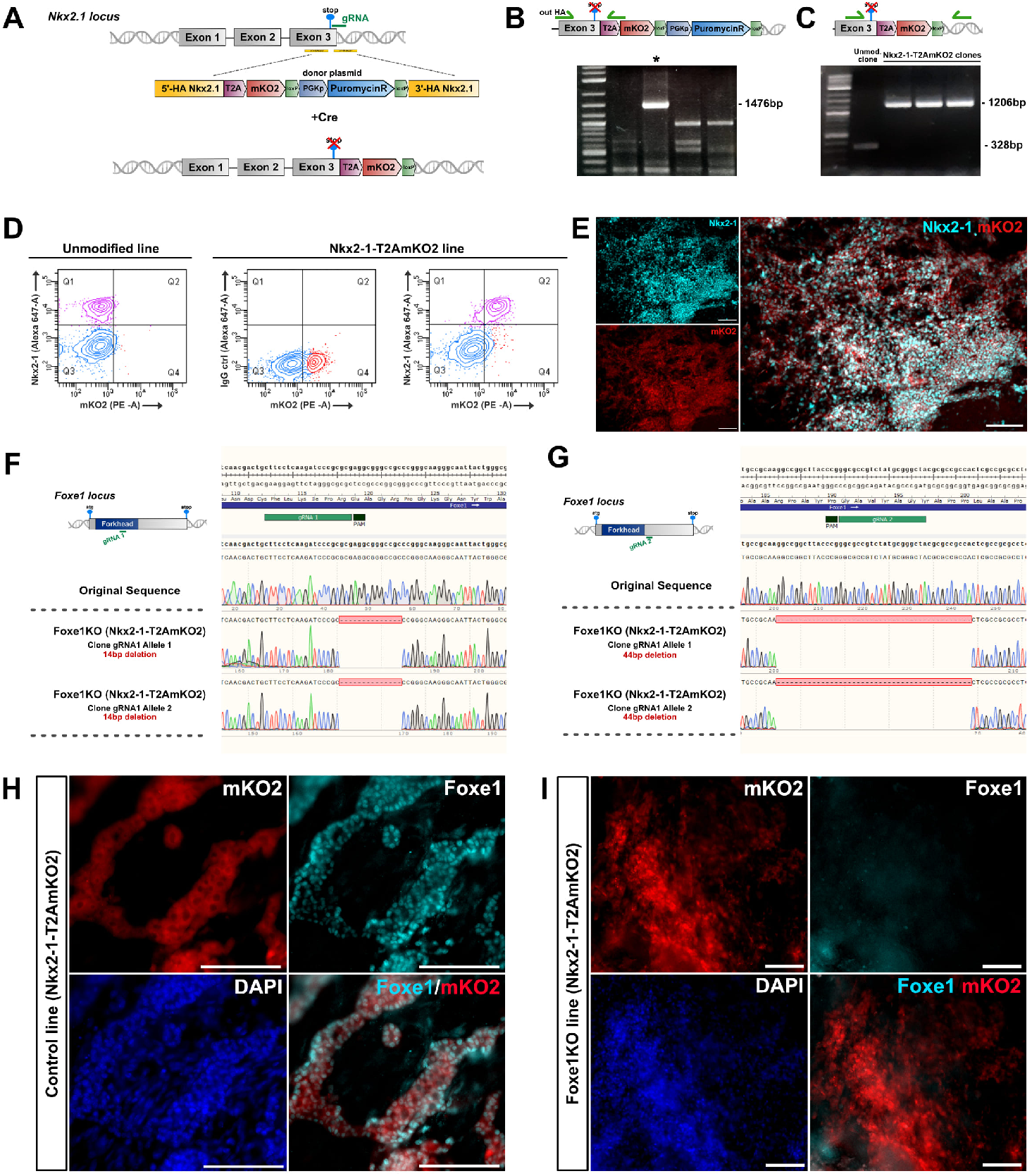
Generation of endogenous Nkx2-1 reporter lines. (**A**) Schematic representation of CRISPR-mediated knock-in strategy to insert T2AmKO2 cassette in 3’-UTR region of Nkx2-1 loci. (**B-C**) PCR screening of knock-in generated clones before (**B**) and after (**C**) PuroR gene excision. Asterisk (*) on image in B indicates a correctly integrated clone. (**D-E**) Validation of the Nkx2-1-T2AmKO2 line. (**D**) Cells from day 14 of thyroid differentiation protocol were subjected to Nkx2-1 immunostaining followed by flow cytometry. Observe a double-positive stained population in Q2, only in the cAMP treated cells derived from Nkx2-1-T2AmKO2 line. (**E**) Images showing Nkx2-1 and mKO2 co-staining at day10 2D culture. (**F-I**) Generation of Foxe1KO/Nkx2-1reporter line. (**F-G**) Genomic profiling of a Foxe1KO/Nkx2-1 reporter clones obtained with guide RNAs targeting inside (**F**) or outside (**G**) the Forkhead domain. (**H-I**) Foxe1 immunostaining of control and Foxe1KO/Nkx2-1 reporter cells subjected to thyroid differentiation. Absence of Foxe1 protein expression is observed in Foxe1KO cells (**I**). Scale bars : 100 µm.

**Figure S5.**
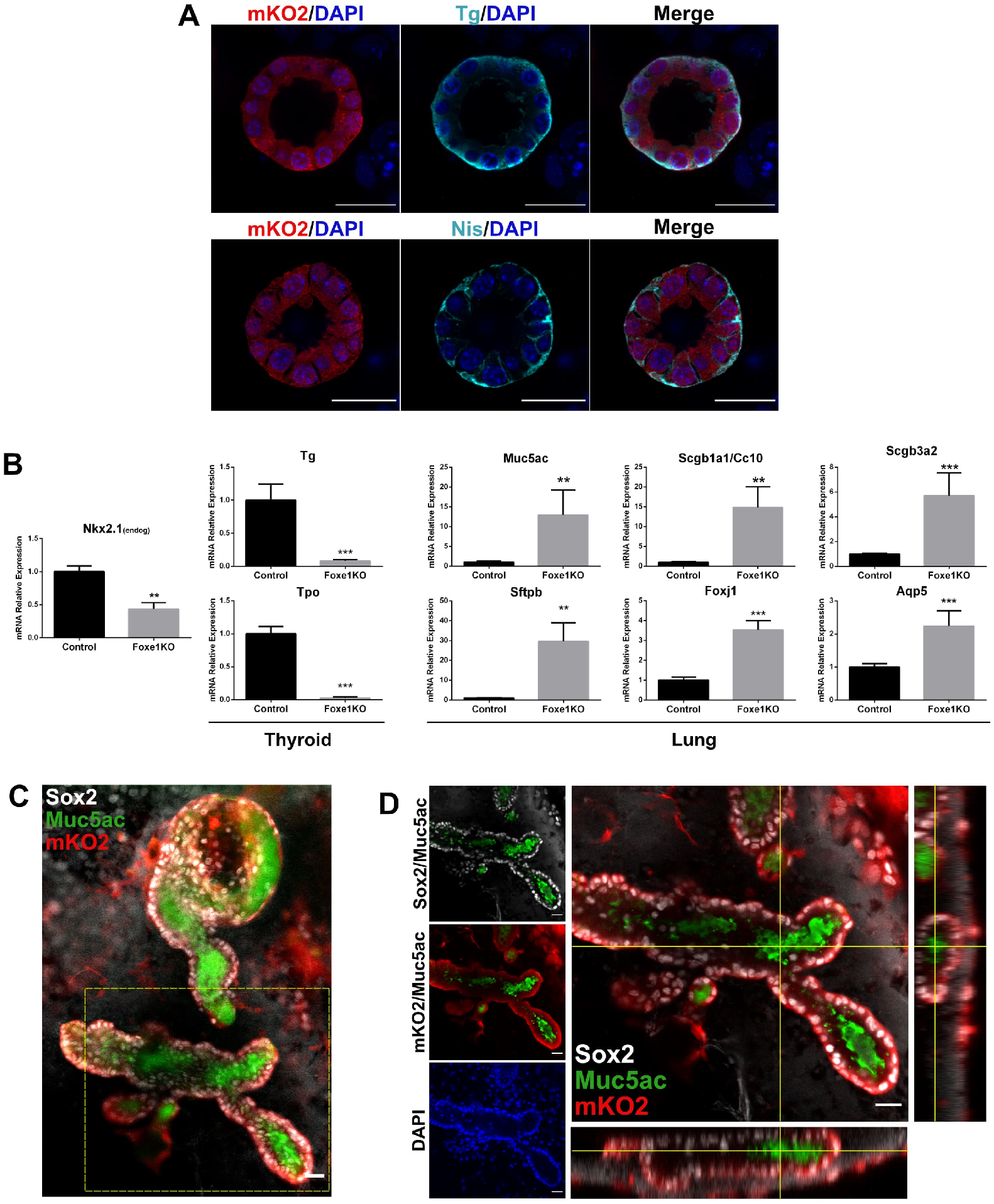
Thyroid and lung differentiation in control and Foxe1KO Nkx2-1 reporter lines. (**A**) Thyroid follicle formation in control Nkx2-1 reporter line observed by co-immunostaining of Tg/mKO2 and Nis/mKO2. (**B**) RT-qPCR analyses of thyroid and lung expressed genes in control and Foxe1KO lines. Observe the reduction of Tg and Tpo thyroid genes whereas lung markers are up-regulated in Foxe1KO cells. (**C-D**) Foxe1KO Nkx2-1 reporter cells differentiation into lung airway organoids. (**C**) Immunostaining for Sox2, mKO2 and Muc5ac. (**D**) Higher magnification and orthogonal views by confocal microscopy of lung organoid depicted in C. Scale bars : 25µm.

**Figure S6.**
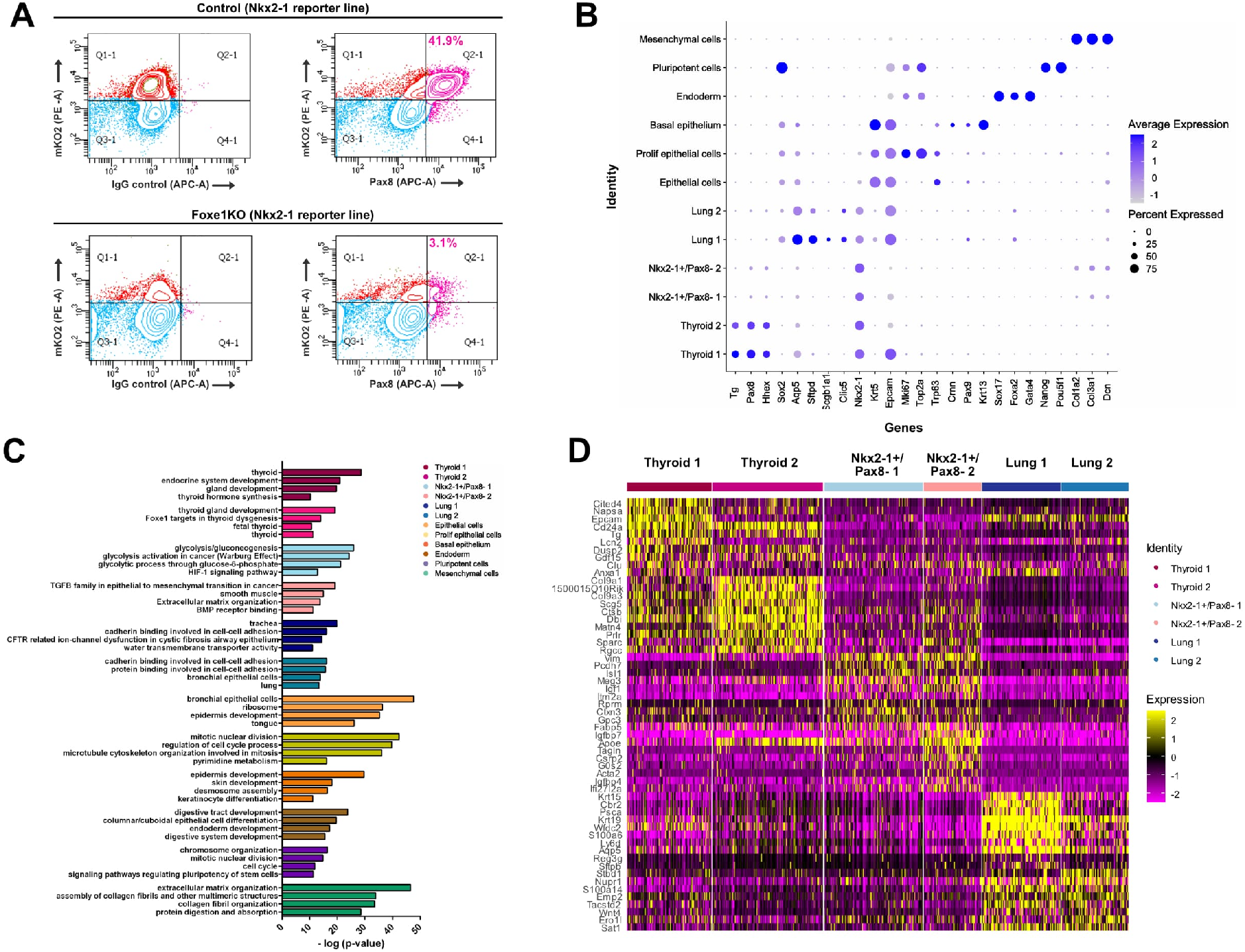
scRNAseq profiling of Foxe1KO cells. (**A**) Pax8 immunostaining followed by flow cytometry analyses at day22. Numbers shown in Q2-1 quadrant in images on the right correspond to percentage of double positive Pax8 (APC) and mKO2 (PE) in control and Foxe1KO cells. Images on left: isotype control. (**B**) Average expression per cluster of selected markers used to define cluster identity. (**C**) Gene Ontology (GO) and pathway analysis performed with top50 differentially expressed genes in each cluster. (**D**) Heatmap of top 50 differentially expressed genes in each cluster expressing Nkx2-1.

### Supplementary Tables

**Table S1.** List of signature genes used in Module Score function and Foxe1 predicted targets (.xlsx file)

**Table S2.**
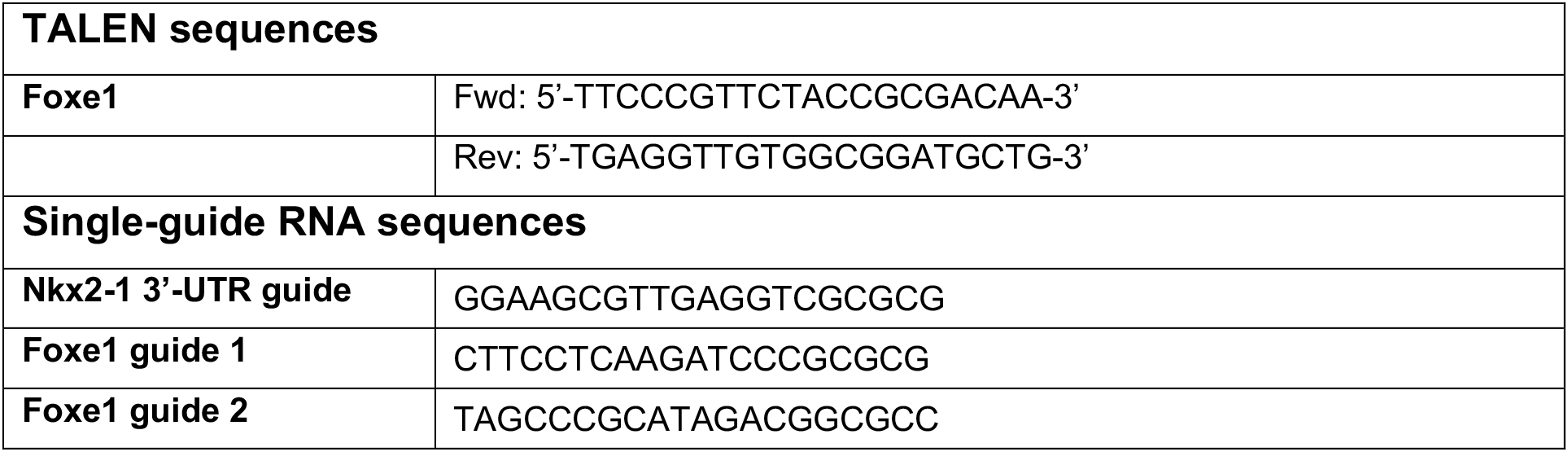
TALEN and single guide RNA sequences.

**Table S3.**
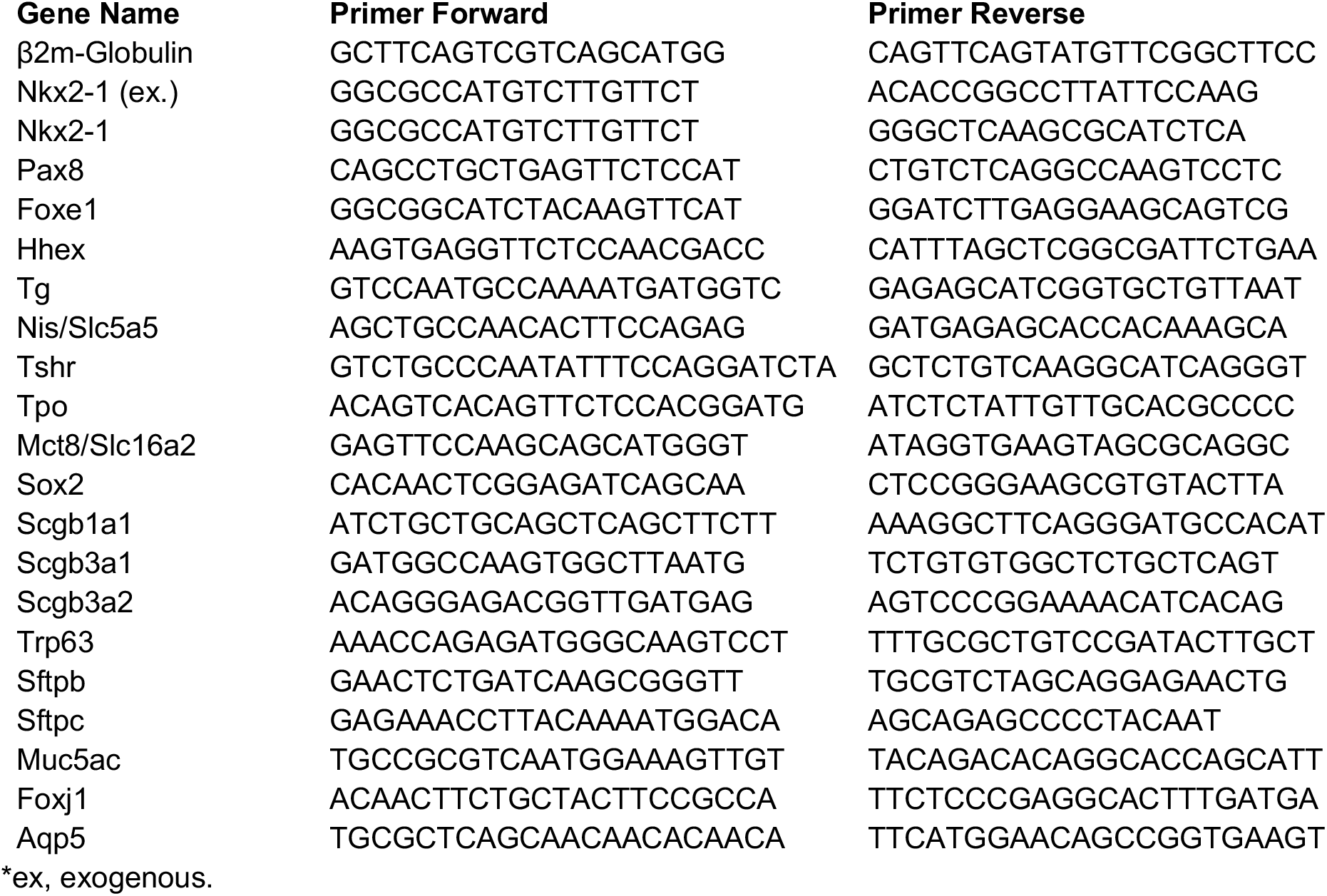
List of primers sequences used for qRT-PCR analysis.

